# CholBindNet: Interpretable Neural Networks for Cholesterol Binding Site Prediction

**DOI:** 10.64898/2026.01.02.697429

**Authors:** Alexis Hernandez, Aashish Bhatt, Ivan Revilla, Jacob Ede Levine, Sai Chandra Kosaraju, Yun Lyan Luo

**Affiliations:** Computer Science Department, California Polytechnic State University, Pomona, USA; Department of Biotechnology and Pharmaceutical Sciences, Western University of Health Sciences, Pomona, USA

## Abstract

Cholesterol is a key modulator of membrane protein structure and function, yet predicting cholesterol binding sites remains challenging due to its undrug-like physicochemical properties. Here, we curated more than 800 high-resolution transmembrane protein structures containing cholesterol and developed an interpretable, atom-based deep-learning framework, CholBindNet, comprising four model architectures: a 3D convolutional neural network, a graph neural network, a graph attention network, and a graph convolutional network. A Positive–Unlabeled (PU) training strategy was employed to address the scarcity of true negative samples resulting from the promiscuous nature of cholesterol binding. We show that CholBindNet substantially outperforms existing deep-learning models trained on general ligand-binding datasets. The performance and generalizability of the model were further demonstrated by rapidly assessing strong, median, and weak cholesterol-binding sites in the PIEZO2 ion channel in excellent agreement with computationally expensive all-atom molecular dynamics (MD) simulations. Additionally, strong model interpretability was achieved for CholBindNet through atom-level feature encoding, Grad-CAM visualization, and attention-based scoring analysis. Overall, CholBindNet provides an efficient and scalable approach for predicting cholesterol binding sites on membrane proteins, achieving performance comparable to MD simulations while offering mechanistic biophysical insights beyond amino-acid sequence. This work hence lays the foundation for future development of deep-learning models targeting membrane protein drug-binding sites and cholesterol-modulated therapeutics.

**Significance Statement:** Deep-learning models for ligand-binding prediction have advanced rapidly, yet those trained on soluble proteins perform poorly for membrane proteins, particularly for cholesterol binding. We introduce CholBindNet, a set of neural network models specifically designed to identify cholesterol-binding sites in transmembrane proteins. CholBindNet substantially outperforms existing deep-learning approaches and accurately ranks strong, intermediate, and weak cholesterol-binding sites in close agreement with computationally intensive all-atom molecular dynamics simulations. This work provides a practical and scalable alternative to long-timescale simulations for studying cholesterol–protein interactions. The curated cholesterol benchmark and open-source models will enable broader adoption of deep learning for investigating lipid regulation of membrane proteins and for guiding the design of drugs targeting membrane-embedded binding sites.

## 1 Introduction

Cholesterol is a vital component of eukaryotic cell membranes, comprising ∼40 mol % of the composition of the plasma membrane [1]. Beyond its role in modulating membrane mechanical properties and sustaining leaflet asymmetry, cholesterol has been shown to modulate many membrane proteins through direct binding [2–6]. Cholesterol is a polycyclic amphipathic molecule, whose polar side consists of a single hydroxyl group that forms hydrogen bonds with lipid headgroups or protein residues. Its apolar section has an asymmetric planar structure, with a rough, aliphatic *β* face and a smooth *α* side. It has long been proposed that branched apolar residues such as Ile, Val, and Leu interpenetrate the rough *β* face of cholesterol, whereas aromatic side chains preferentially stack against its planar *α* face [2]. Although physically plausible, this heuristic interpretation alone is insufficient to account for the wide diversity of cholesterol-binding pocket topologies observed across membrane proteins.

Experimental techniques used to identify cholesterol binding sites, including X-ray crystallography, cryo-electron microscopy (cryo-EM), NMR, and fluorescent labeling, are resource-intensive and time-consuming [6]. Thus, developing reliable and efficient computational approaches for predicting cholesterol binding is highly desirable. Nguyen and Ondrus [7] provided a comprehensive review of in silico tools developed to predict cholesterol-binding sites, noting that traditional bioinformatics approaches using sequence motifs (CRAC, CARC, CCM) often yield high false-positive rates due to their permissive patterning. More refined strategies, such as template-based tools like CholMine utilize feature-matching to known binding sites [8] and RosettaCholesterol, leverage a database of 55 then-available cholesterol-protein structures to improve the docking accuracy [9]. However, the efficacy of these methods is limited by the small data set. A more broadly applicable approach is molecular dynamics (MD) simulations, in which cholesterol binding/unbinding events can be monitored and ranked from simulations of a protein embedded in a cholesterol-containing bilayer. To achieve microseconds of MD simulations, less accurate coarse-grained models, such as the eight-particle Martini model of cholesterol, are usually used instead of atomistic models [10]. However, MD simulation is not yet suitable for high-throughput prediction of cholesterol binding across many protein targets.

Recent years, advances in computer technology and the rapid expansion of protein structure databases have fueled the development of machine learning models for the prediction of protein-ligand binding. Neural networks have become increasingly valuable for predicting drug binding sites, thanks to their ability to learn intricate, nonlinear patterns from large-scale biological datasets. These models often outperform traditional computational approaches, offering new possibilities in structural biology and drug discovery [11–20]. However, to the best of our knowledge, no dedicated deep-learning models have been developed to predict cholesterol binding sites. We show in this study that existing deep learning models, Chai-1 [17], DiffDock [16], LABind [18], trained primarily on druglike ligands, are not directly applicable to cholesterol. Cholesterol binding sites often exhibit unique structural and physicochemical characteristics, such as proximity to membrane-facing surfaces, and hydrophobicity, which may be overlooked by generic binding site predictors [7]. In addition, generic binding site predictors often struggle to predict shallow binding pockets [11], which is a prominent feature of binding sites that reside in the transmembrane region.

Recent advances in studying membrane proteins in their native environments have provided a rapidly expanding collection of cholesterol-binding sites across major classes of drug targets, including G protein–coupled receptors, ion channels, transporters, and nuclear receptors [21, 22]. We curated more than 800 high-resolution X-ray crystallographic and cryo-EM structures of transmembrane proteins with bound cholesterol from the RCSB PDB. Using experimentally observed binding sites as positive data and docking-generated sites as unlabeled data, we applied our recently developed Positive–Unlabeled (PU) learning strategy to train four neural network models: a 3D convolutional neural network (Chol-3DCNN), a graph neural network (Chol-GNN), a graph attention network (Chol-GAT), and a graph convolutional network (Chol-GCN). We found that our models not only outperform existing deep-learning approaches in predicting experimental cholesterol-binding sites, but also has the ability to rank strong, medium, and weak cholesterol-binding sites on the mechanosensitive ion channel PIEZO2, in good agreement with all-atom molecular dynamics simulations. Importantly, the interpretability of our models offers rich biophysical insight into the atomic-level spatial patterns underlying cholesterol recognition.

## 2 Results

Our proposed *CholBindNet* framework is an interpretable deep learning model developed to identify cholesterol-binding pockets within membrane proteins. As illustrated in **Figure 1**, it comprises four major components: (A) data preprocessing, (B) deep learning architecture, (C) positive-unlabeled (PU) learning, and (D) interpretability. In the preprocessing stage, experimentally validated cholesterol-binding sites are extracted from the Protein Data Bank (PDB). Each protein is represented as a one-hot encoded matrix integrated with structural distance information to capture spatial context (**Figure 1A**). This representation serves as input to the deep learning pipeline, which integrates graph-based neural networks (Chol-GNN), graph convolutional neural network (Chol-GCN), graph attention mechanisms (Chol-GAT), and three-dimensional convolutional layers (Chol-3DCNN) to learn hierarchical spatial and chemical features (**Figure 1B**). The output layer of deep learning architecture produces probabilistic scores for cholesterol-binding likelihood under a PU-learning objective, enabling robust prediction even with limited positive labels (**Figure 1C**). Finally, an interpretation module (**Figure 1D**) quantifies feature-level contributions and reveals previously uncharacterized binding phenomena, providing mechanistic insights into cholesterol–protein interactions.

**Figure 1.**
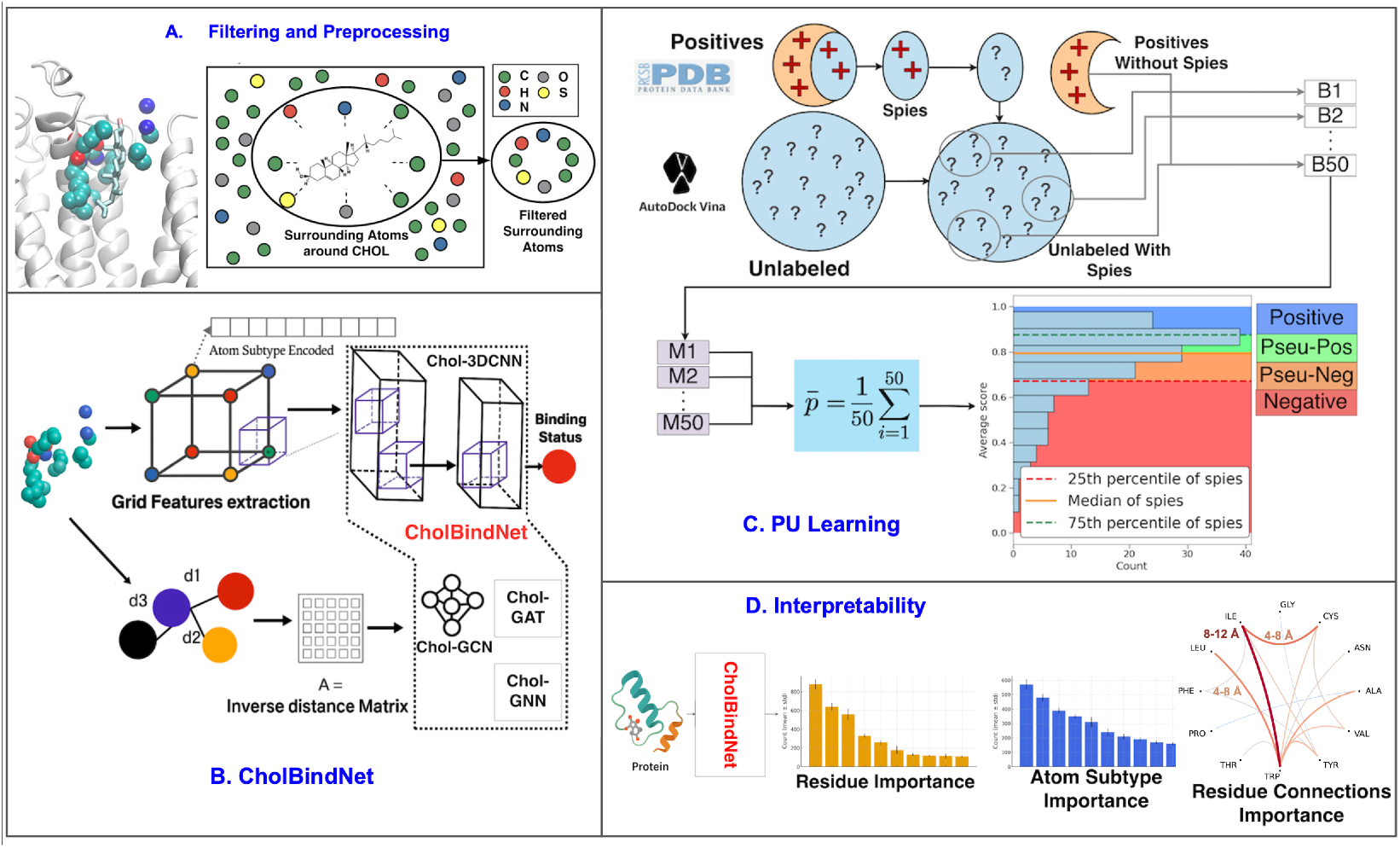
Positive-unlabeled deep learning framework for predicting cholesterol binding sites. (A) Filtering and Preprocessing: Only atoms within 5 Å of the cholesterol ligand are retained, and each atom subtype is one-hot encoded to generate the input representation. (B) CholBindNet Architecture: CholBindNet integrates four model architectures, with the CholBind Graph Neural Network (Chol-GNN) variant achieving the highest accuracy across multiple datasets. (C) Positive–Unlabeled (PU) Learning Strategy: To address data imbalance and enable robust evaluation, PU learning is employed with “spy” samples; known positive samples temporarily relabeled as unlabeled. This allows the model to estimate its ability to recover hidden positives. The dataset is divided into 50 bins (B1–B50), and a separate submodel (M1–M50) is trained for each bin. Predictions from the 50 submodels are then averaged for stable performance. Spy samples are evaluated to obtain probability distributions, and thresholds based on the 25th, 50th, and 75th percentiles of spy probabilities are used for relabeling with 4 possible labels: Negative, Pseudo-Negative, Pseudo-Positive, and Positive. This process is termed as percentile labeling. (D) Model Interpretability: Post-processing methods are used to identify the most important residues, atom subtypes, and residue–residue connections contributing to cholesterol binding.

### 2.1 Positive and Unlabeled Data Sets

We extracted 770 high-resolution (≤ 3 Å) X-ray crystallographic or cryo-EM structures of membrane proteins containing cholesterol from the RCSB Protein Data Bank Database (until January 2025) as positive samples. We excluded cholesterol hemisuccinate in our samples, which is a solubilized derivative of cholesterol used during protein sample preparation. Because no negative data have been reported in the literature, and cholesterol is likely to bind to multiple locations within a single protein, we generated unlabeled data by performing global docking of cholesterol molecules into each protein from the positive data. Because AutoDock Vina scoring function was not benchmarked on cholesterol data set, we expect it to generate both true and false positive cholesterol binding sites. Around four distinct binding sites per protein from Vina docking are saved as unlabeled data. This approach is advantageous over assigning randomly selected atoms outside known binding sites (e.g., atoms beyond 5 Å of any ligand atom) as negatives, because the latter is unlikely to reflect genuine binding pocket features and may compromise the model’s ability to detect false positives. The total cholesterol data set consists of 770 positive and 2762 unlabeled samples.

### 2.2 Positive-Unlabeled (PU)-learning data splits

For PU-learning, we randomly chose 20% of the positive data to be spies. These spies are trained as unlabeled [23]. We did this to check if the models can accurately identify positives that were labeled as unlabeled. The probability of identifying the spies as positive is then used for classification (i.e., percentile labeling). The rest of the positive samples are split into 50/10/20% as training/validation/test sets.

Because Vina docking was conducted on positive sample proteins without cholesterol present in the binding site, some of the top binding sites from global docking overlap with the positive binding sites. We hence moved 369 unlabeled samples that overlapped with positive binding sites to the unlabeled test set to evaluate models’ ability to distinguish positives from unlabeled samples. The remaining unlabeled samples (2762 - 369 = 2393) were then divided into five independent experiments, each using a distinct data split configuration as outlined in **Figure 2**.

**Figure 2.**
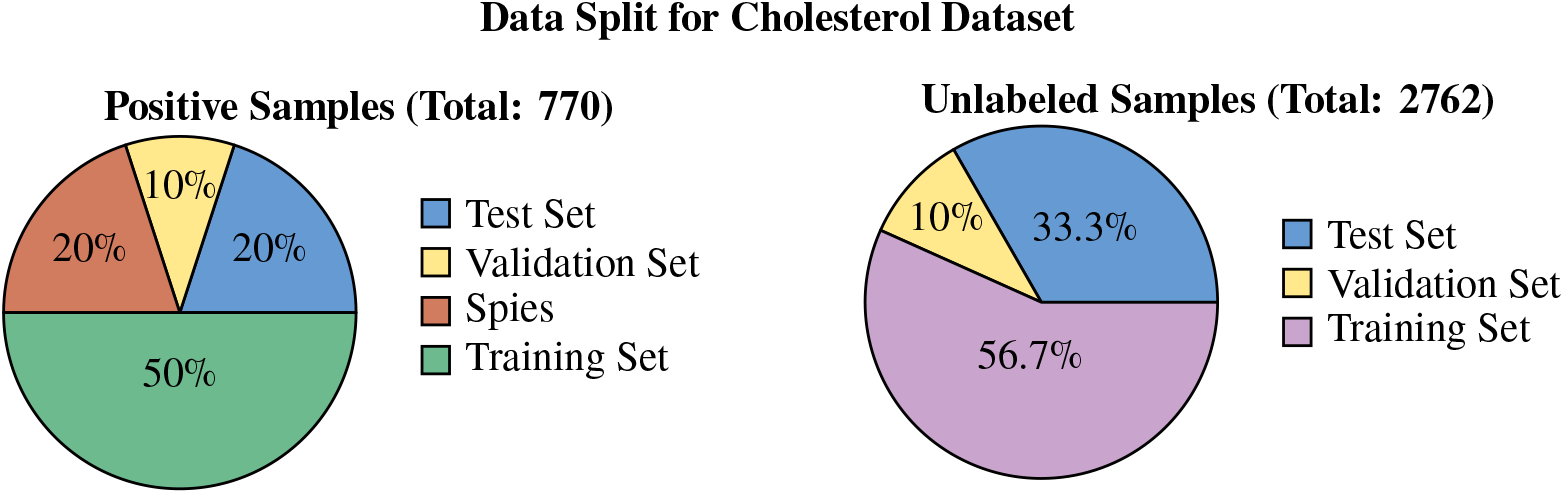
Data split for cholesterol dataset. The same process was applied to all 5 experiments with different randomly chosen samples for each group.

### 2.3 External validation and PIEZO2-cholesterol binding sites

In addition to the positive and unlabeled samples, we included two independent datasets to evaluate the generalization of our model predictions. The first is a set of 57 additional cholesterol bound cryo-EM structures that we obtained from our in-house query script that captured more transmembrane proteins than the web-based query. Those protein structures were not used to generate unlabeled data and were excluded from training and validation. We call it the external validation set.

The second is 109 strong or weak binding sites generated from all-atom molecular dynamics (MD) simulations of a mechanosensitive PIEZO2 ion channel in a membrane bilayer model containing 1-palmitoyl-2-oleoyl-glycero-3-phosphocholine (POPC) and cholesterol. PIEZO channels are one of the largest plasma transmembrane proteins known to date, each consists of 114 transmembrane helices. Although cholesterol has been reported to modulate or being modulatd by PIEZO channel function [24–27], experimental cholesterol binding sites of the PIEZO2 channel are not known. Using two replicas of a total of 28 *μ*s MD trajectories generated on the Anton3 supercomputer, we ranked each binding site according to the percentage of cholesterol occupancy over the entire trajectory. As it is difficult to capture negative binding sites from MD simulation, we also supplemented 23 binding sites from Vina global docking that showed only 10-20 % occupancy from MD simulation.

### 2.4 Data Preprocessing

Protein atoms within 5 Å of cholesterol were selected as binding sites in both positive and unlabeled samples. After collecting these surrounding atoms, we one-hot encode the subtype of each atom (e.g., C, CA, CB) (**Table 1**). These atom subtypes contain information on the type of atom, as well as its location on each amino acid. For atom *i*, let **x**_*i*_ ∈ {0, 1}^37^ denote its one-hot vector; stacking rows gives the feature matrix **X**. All of our models share this atom-subtype encoding, which allows for direct comparisons between architectures.

**Table 1.** Shared Input Atom Subtypes (37 One-Hot Channels)

| Atom Class | Subtypes |
| --- | --- |
| Carbon (17) | C, CA, CB, CD, CD1, CD2, CE, CE1, CE2, CE3, CG, CG1, CG2, CH2, CZ, CZ2, CZ3 |
| Oxygen (8) | O, OH, OD1, OD2, OE1, OE2, OG, OG1 |
| Nitrogen (9) | N, NE, NE1, NE2, ND1, ND2, NZ, NH1, NH2 |
| Sulfur (2) | SD, SG |
| Other (1) | UNKNOWN |

### 2.5 PU Learning

The PU learning strategy shown in **Figure 1C** enabled model training in the absence of reliable negative labels by constructing an ensemble of balanced sub models from the training split. Each sub model paired positive training samples with an equal number of unlabeled samples drawn from a larger unlabeled pool and treated as weak negatives, with different unlabeled subsets contributing across the ensemble to capture variability within the unlabeled space. Averaging probability scores across the 50 sub models for each architecture produced consensus predictions that were less influenced by individual model behavior or by specific negative assignments (see methods).

The final assignments to the class were determined using a custom *percentile labeling* (as illustrated in algorithm 1) based on the empirical distribution of the spy scores rather than fixed probability cutoffs. By assigning labels according to the relative score ranking in the 25th, 50th and 75th percentiles, the classification reflected the structure of the learned score distribution and facilitated the consistent separation of samples across architectures. The comparative results for alternative thresholding strategies, including a conventional binary cutoff, are presented in **Figures S1– S4**. In the context of PU learning, these results indicate that conventional fixed probability thresholds do not adequately reflect the underlying score distribution, while percentile labeling offers a more appropriate distribution-aware mechanism for assigning class labels. We also note that the Chol-GNN and Chol-GCN class assignments are consistent across different percentile labeling, demonstrating the robustness of the models.

### 2.6 Models Performance

Model performance was evaluated using five independent experimental runs, each implementing the full PU-learning procedure with 50 sub models per architecture. Evaluation on the positive and unlabeled test sets, as well as on an external validation cohort, showed consistent separation of unlabeled samples from the positive class (**Figure 3**). The resulting class distributions indicate that graph-based models capture structural features associated with cholesterol binding, particularly for binding sites characterized by compact geometries and recurring hydrophobic interaction patterns.

**Figure 3.**
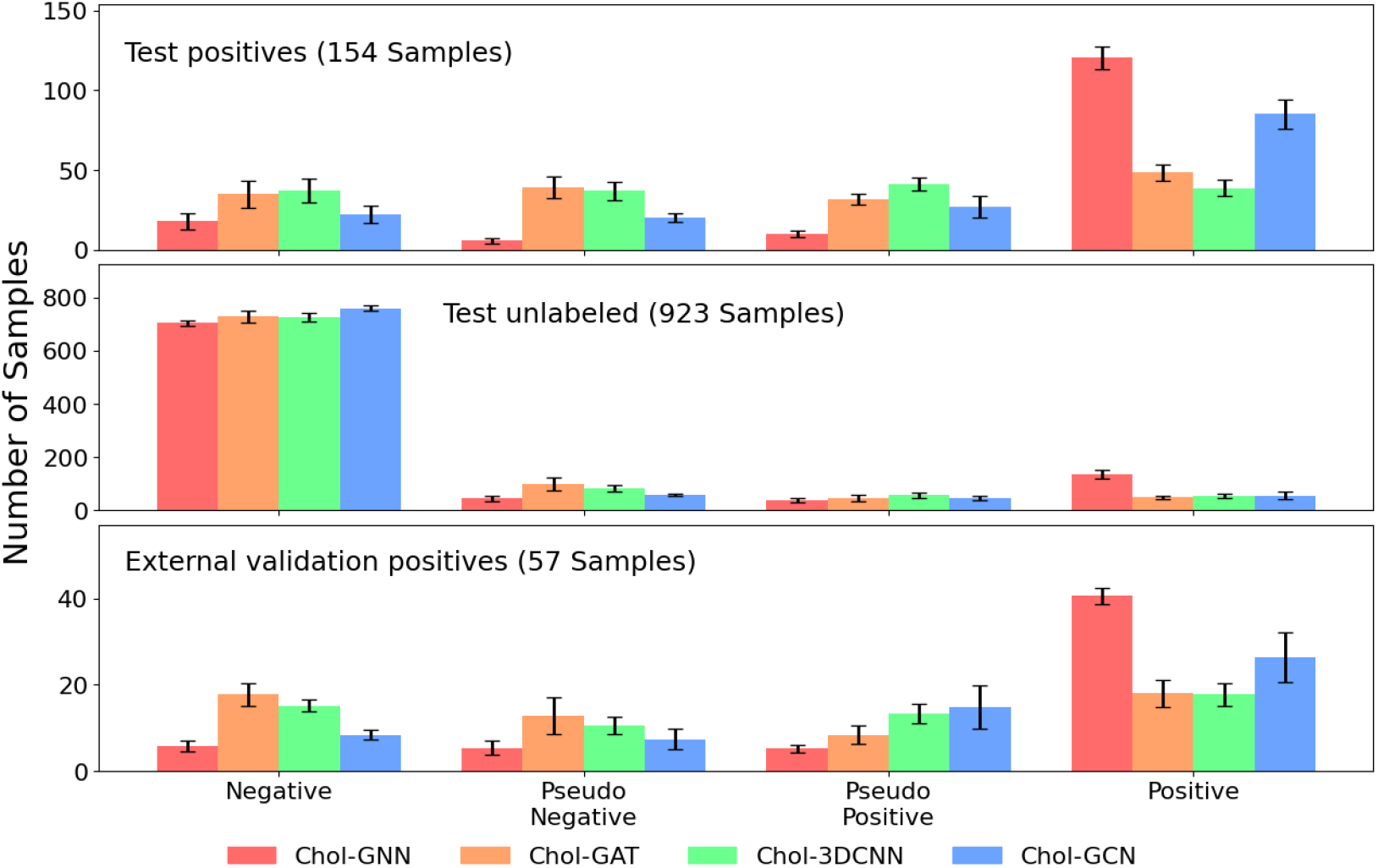
Performance of the percentile evaluation method. Results are shown for the test positive set, test unlabeled set, and external validation positives. Each bar represents the mean performance across five independent experiments, with error bars indicating the standard deviation. The percentile evaluation method applies labeling thresholds based on the 25th, 50th, and 75th percentiles of spy-sample probability distributions.

All graph-based architectures operate on identical pocket-derived node features and adjacency representations, and thus encode the same underlying physicochemical information. Differences in predictive behavior therefore primarily reflect how each architecture aggregates and propagates these features. The attention mechanism employed in Chol-GAT substantially increases model capacity, but for cholesterol-binding sites with relatively constrained pocket sizes and limited interaction heterogeneity, this added expressiveness does not appear necessary to resolve relevant binding determinants. In such settings, simpler message-passing formulations are sufficient to capture the structural regularities governing cholesterol association. This interpretation is further supported by the observation that Chol-GAT exhibits comparable behavior to the other graph-based models in the PIEZO2–cholesterol binding site ranking task (Section 2.9), which involves greater structural diversity and interaction complexity. Together, these results suggest that the effectiveness of increased model capacity depends on the complexity of the binding-site landscape rather than on architectural sophistication alone.

The spy score distributions produced by the Chol-3DCNN were consistently concentrated near the upper end of the probability range across all five runs (**Figure S4**). Median spy scores exceeded 0.997, and even the lower quartiles remained high, indicating limited variability in predicted probabilities. A similar pattern was observed when classical atom-level features computed using RDKit were used to train the 3DCNN model (**Figure S5**), suggesting that this behavior is intrinsic to the architecture rather than to the feature representation. The full list of RDKit features is provided in **Table S1**.

Within the unlabeled test set, all models assigned a small subset of samples to positive or pseudo-positive classes. To assess whether these predictions correspond to sites resembling known positives, we computed an overlap score defined as the number of atoms shared between an unlabeled site and a known positive site. As shown in **Figure 4**, samples classified as positive, pseudo-positive, or pseudo-negative exhibit higher average overlap scores than those classified as negative, indicating that the models identify unlabeled sites that are structurally closer to known cholesterol-binding pockets.

**Figure 4.**
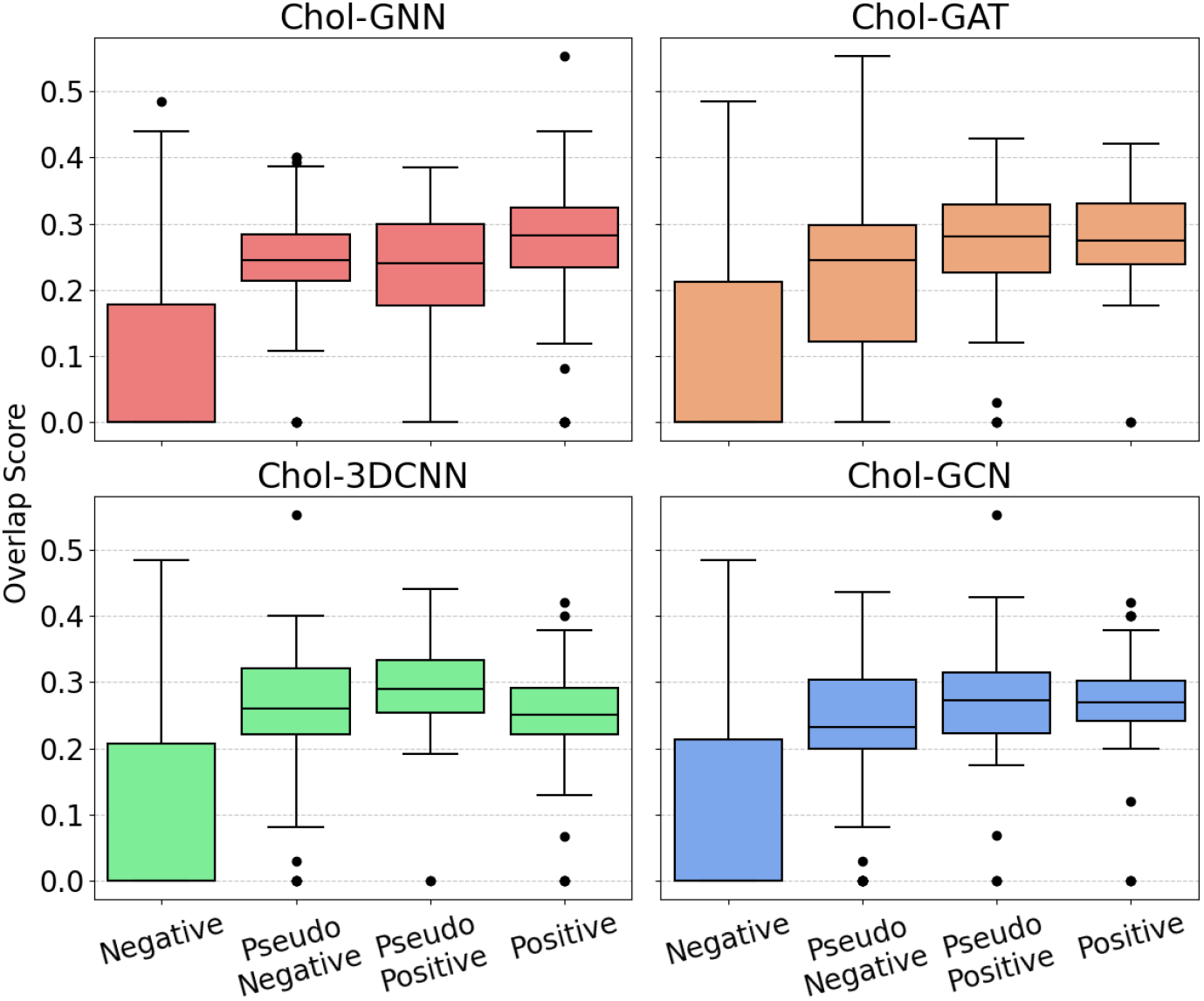
Overlap scores of the binding sites between unlabeled test set and positive samples. Overlap scores are used to quantify how closely an unlabeled binding site aligns with a known positive cholesterol binding site across four percentile-based classification groups: Negative, PseudoNegative, PseudoPositive, and Positive. Box plots are shown for each of the four model architectures (Chol-GNN, Chol-GAT, Chol-3DCNN, and Chol-GCN).

Model performance was further examined to assess potential dependence on binding-pocket atom count. **Figure S6** shows the relationship between predicted probability scores and the number of atoms retained after filtering surrounding residues around cholesterol. For samples containing approximately 20 to 60 atoms, positive and unlabeled cases span a similar range of predicted scores, with no evident shift favoring either class. This pattern is consistent with predictions that are not strongly influenced by atom count within this range. In contrast, samples with larger atom counts are predominantly associated with low-scoring unlabeled cases, and true positives are rarely observed beyond approximately 90 atoms. This is likely because Vina docking tends to position cholesterol molecules deeper within the transmembrane region without accounting for binding-site accessibility or the orientation of cholesterol relative to the membrane bilayer.

### 2.7 Model Interpretation

In this study, our goal is not only to leverage deep learning to predict cholesterol-binding sites, but also to uncover the hidden patterns that underlie their biophysical properties. The cholesterol recognition amino acid consensus (CRAC) motif has been the most widely discussed cholesterol-binding domain in the literature for more than two decades. CRAC, along with its inverted counterpart CARC, represents a bidirectional linear motif characterized by a branched apolar residue (Leu or Val), followed by a stretch of one to five residues of any type, an aromatic residue (Tyr or Phe), another flexible segment of one to five residues, and finally a basic residue (Arg or Lys). The looseness of this motif definition suggests that cholesterol binding is not strictly sequence-specific, but rather structurally promiscuous, relying more on the local three-dimensional environment than on any fixed residue pattern. Our models are designed to capture this three-dimensional structural context directly from atomic coordinates, rather than relying on one-dimensional sequence motifs. Notice that we deliberately did not categorize our data set based on whether cholesterol binds at the protein–membrane interface or between transmembrane (TM) helices, as such a classification can introduce unwanted bias (e.g., many cholesterols that bind between TM helices are still exposed to lipids).

#### 2.7.1 Residue importance

All of our models are atom-based, using only atom subtypes and spatial coordinates as node and edge (or grid) features. No preselected chemical descriptors or residue types are provided as input features. Consequently, in our model interpretation, residue importance is inferred solely through back-mapping from atomic features to their corresponding residues. Remarkably, the residue-level importance rankings obtained in this way are highly consistent across the Chol-3DCNN, Chol-GNN, and Chol-GCN architectures, with branched apolar residues (Leu, Ile, Val) and aromatic residues (Phe, Trp) ranked as the most influential for cholesterol binding (**Figure 5**). Interestingly, the Chol-GAT model assigns the highest importance to Trp among all residue types. Analysis of residue pairs importance suggests that it is due to the GAT emphasizing long-range interactions (see section 2.7.3 below). Although these residue importance trends are overall consistent with the CRAC motif definition, our atom-level importance analysis reveals that only a small subset of atoms within these residues predominantly contribute to cholesterol binding, and their spatial arrangement is important.

**Figure 5.**
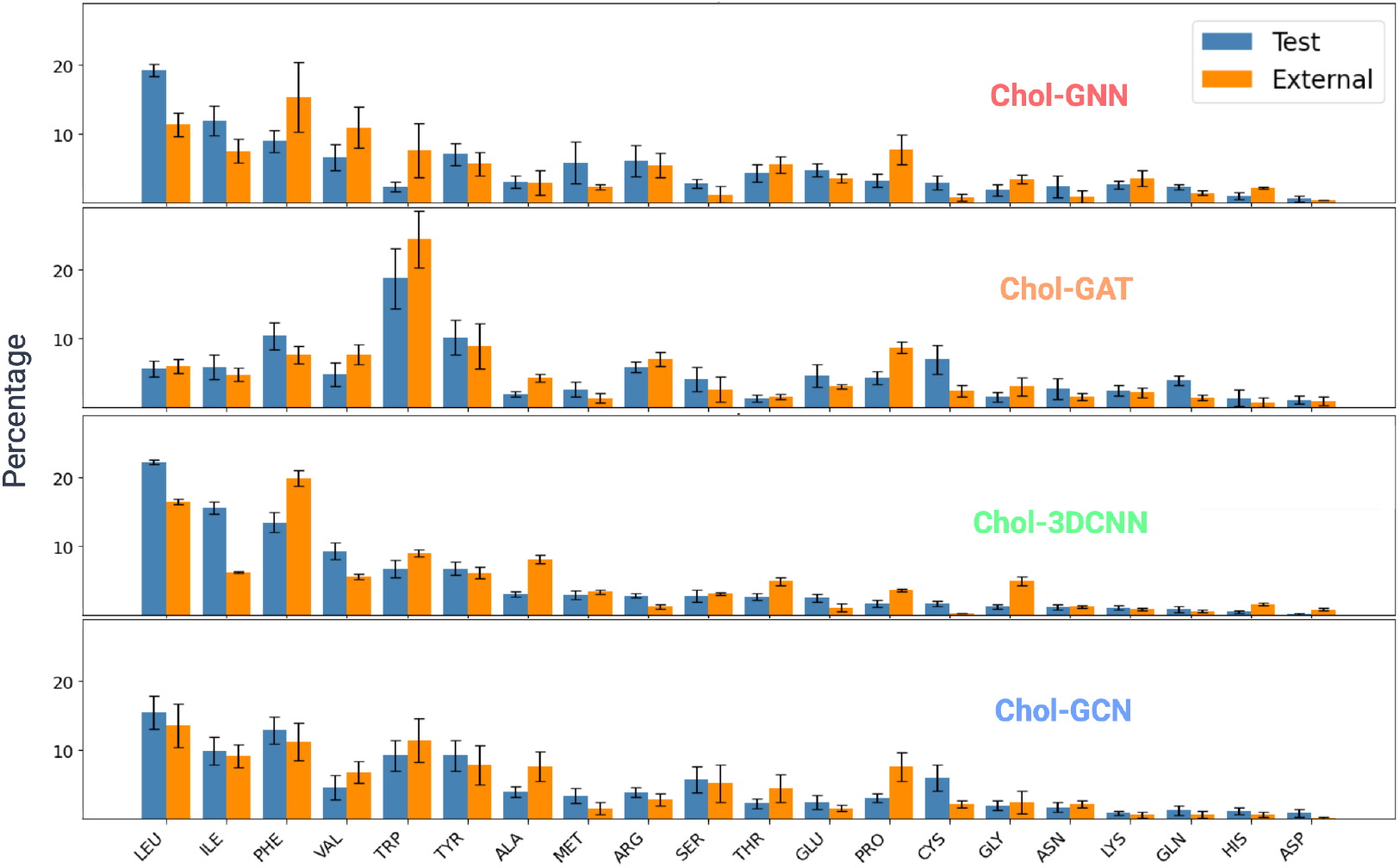
Residue importance in predicting cholesterol binding sites. The bar plots show the mean normalized frequency of each residue ranked among the top 5 most important features in each model, for the test positive set and external validation positive datasets. Error bars indicate the standard deviation across five experiments.

#### 2.7.2 Atom importance

At the atomic level, the most influential atom subtypes include backbone atoms (C, O, CA), common side-chain carbons (CB, CG), and the CD1, CD2, and CZ atoms found in aromatic residues (Phe, Tyr, Trp) and in Arg, as well as CG2 atoms located on the branched aliphatic residues Ile, Thr, and Val (**Figure 6**). These results suggest that backbone carbonyl oxygen atoms, rather than side-chain heteroatoms, form the majority of hydrogen-bonding interactions with cholesterol’s polar 3-hydroxyl group, which typically orients toward the membrane surface [2]. A small subset of carbon atoms, predominantly hydrophobic or aromatic, contributes the strongest interactions with cholesterol’s rigid, nonpolar tetracyclic ring system. Aliphatic side chains such as those of Leu, Ile, and Val provide extensive van der Waals contacts, enabling tight packing against the sterol rings and stabilizing cholesterol within the transmembrane environment. Aromatic residues, including Phe, Tyr, and Trp, engage in CH–*π* interactions with cholesterol’s rings, offering additional stabilization and orientational specificity.

**Figure 6.**
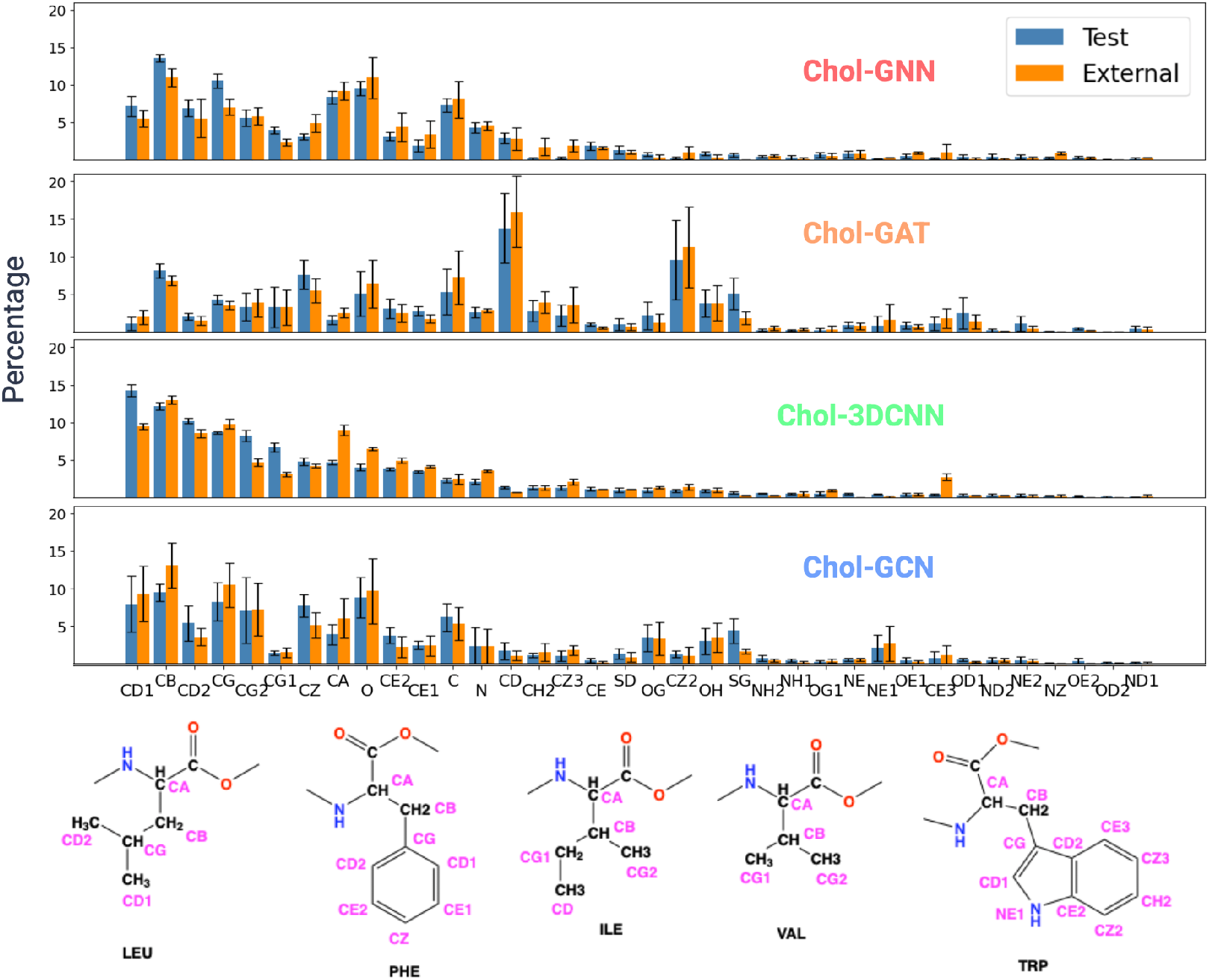
Atom importance in predicting cholesterol binding sites. The bar plots show the mean normalized frequency of each atom subtype ranked among the top 5 most important features in each model, for the test positive set and external validation positive datasets. Error bars indicate the standard deviation across five experiments. Bottom diagrams are examples of atom subtypes in important residues.

#### 2.7.3 Residue pair importance

Although our models provide consistent rankings of important residue and atom subtypes, we hypothesize that the spatial arrangement of these atoms also plays a critical role in cholesterol binding.

To examine this, we leveraged the models’ ability to capture three-dimensional structural context directly from atomic coordinates and extracted important atom–atom and residue–residue pairs across three distance ranges: 4–8, 8–12, and 12–16 Å.

In **Figure 7**, we compare the top 20 most important residue pairs, averaged over all positive test set and external validation samples. In the GAT model, the Ile–Trp pair at an 8–12 Å separation emerges as the most important, consistent with GAT’s emphasis on long-range interactions and its high importance score for Trp. In contrast, the GCN model identifies only residue pairs within the 4–8 Å range among its top 20, with branched aliphatic pairs such as Leu–Val ranked highest, followed by Leu–Ile.

**Figure 7.**
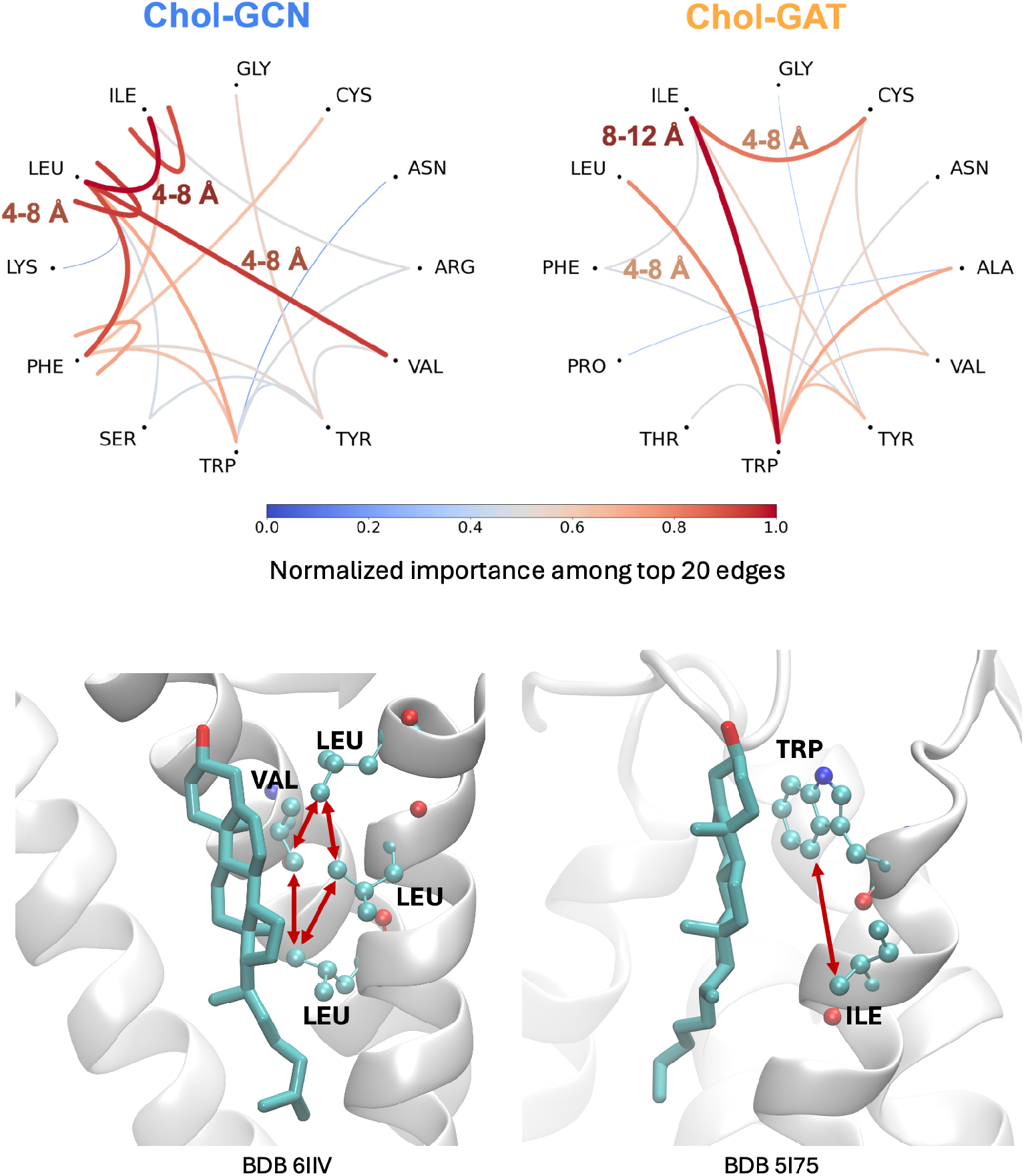
Chord diagram and structural visualization of the most important edges identified by Chol-GCN and Chol-GAT. **Top:** Chord diagrams for Chol-GCN (left) and Chol-GAT (right) illustrate the twenty highest-ranked edges connecting residue pairs within predicted cholesterol-binding pockets. Edge importance is quantified using a combined metric incorporating both frequency of occurrence across experiments and the model’s edge importance score, visualized using a blue-to-red gradient (blue = lower importance; red = higher importance). Distance annotations (4–8 Å, 8–12 Å) highlight the spatial ranges in which the most informative edges commonly appear. **Bottom:** Structural examples rendered in VMD show residue–residue connections corresponding to highly ranked edges detected by the models. The left panel (PDB: 6IIV) illustrates representative Chol-GCN interactions among LEU and VAL residues surrounding the cholesterol molecule. The right panel (PDB: 5I75) shows a prominent Chol-GAT interaction between TRP and ILE near the bound cholesterol.

These atom and residue patterns suggest that cholesterol binding is governed by hydrophobic complementarity and ring stacking. In **Figure 7** bottom, we illustrated the top residue-pair features captured from Chol-CGN and Chol-GAT models. Consistent with previous literature [2], the rough, aliphatic *β* face interchelates with multiple branched apolar residues such as Leu and Val sidechains, and the planar *α* side stacks against aromatic side chains of Trp with an Ile located 9 Å below providing additional hydrophobic interaction stabilizing the other end of the sterol ring. These findings reinforce the biological relevance of the learned model importance patterns and demonstrate that the networks successfully recovered key structural determinants characteristic of cholesterol-binding motifs observed in experimental membrane protein structures.

### 2.8 Model comparison with general ligand binding site prediction methods

To the best of our knowledge, no deep-learning model has been specifically developed to predict cholesterol binding sites. We therefore compared our model’s performance with three classes of deep-learning models designed for general ligand binding prediction: DiffDock [16], Chai-1 [17], and LABind [18]. The comparison was performed using our external validation set, which consists of 57 cholesterol-bound protein structures that were not used during training.

DiffDock successfully predicted cholesterol binding sites within 5 Å of the native binding site for three structures, resulting in a performance score of 5.3% (**Figure 8**). Due to limited GPU memory, Chai-1 was run on a subset of 23 structures containing less than 1000 residues from the external validation set. Among these, Chai-1 correctly predicted the cholesterol binding site within 5 Å of the native binding site for three structures as well, corresponding to a performance score of 13%.

**Figure 8.**
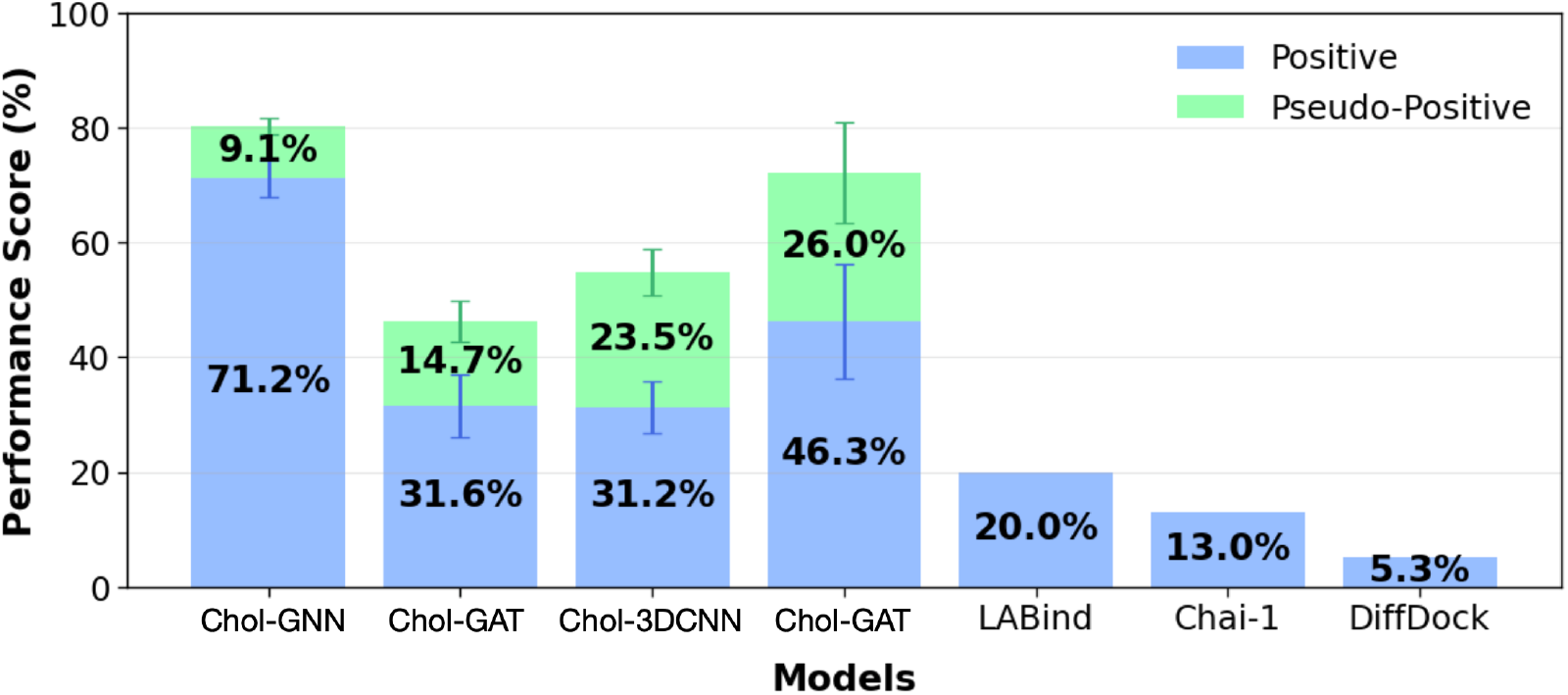
Performance of different models on the external validation dataset. Performance scores represent the proportion of the correctly predicted PDB structures relative to the total number evlauated for each model. LABind was assessed on five structures, Chai-1 on 23 structures and DiffDock on the full set of 57 PDB structures. All three models produced identical results across five repeated runs, hence the absence of error bars. The mean and standard deviation for Chol-GNN, Chol-GAT, Chol-3DCNN, and Chol-GCN are the same as in Figure 3.

Because LABind was designed to handle individual protein chains, only structures in which a single chain contributed to cholesterol binding were suitable for evaluation. However, cholesterol frequently binds at multimeric interfaces, rendering most structures incompatible with LABind. Only five protein chains met this comparison criteria. For these chains, LABind predicted six residues with prediction scores greater than 0.48 for cholesterol binding. Only one of six residues corresponds to a true native binding residue, defined as any residue experimentally observed within 5 Å of cholesterol. Thus, LABind identified the correct binding residues for one of the five evaluated structures, yielding a performance score of 20.0%. Detailed results from DiffDock, Chai-1, and LABind can be found in **Figure S7-S9**. The modest success rate across these three models highlights the specific challenge of predicting cholesterol binding and the need to develop a cholesterol-specific deep-learning model.

### 2.9 Model comparison with MD simulation of PIEZO2 ion channel

To evaluate the robustness and generalizability of our models, we assessed their performance on one of the largest plasma membrane proteins, the PIEZO2 ion channel. Cholesterol modulation of PIEZO1 and PIEZO2 channel functions has been extensively studied [24–27], however, no experimentally resolved cholesterol binding sites have been reported for PIEZO1 or PIEZO2. Here, we used all-atom MD simulations to rank potential cholesterol binding sites on an open state of PIEZO2 channel, by computing the average cholesterol occupancy across two independent MD replicas of 28 *μ*s in total. For each model, we supplied MD snapshots of 109 binding sites with cholesterol occupancy ranging between 20-100%. In addition, we supplied 23 binding sites generated from Vina global docking that showed only 10-20 % occupany during MD simulations.

As shown in **Figure 9** top panel, with the exception of Chol-3D CNN model, the categorical ranking across 132 binding sites is overall consistent with MD-predicted cholesterol occupancy. Among three graphic-based models, Chol-GNN model demonstrated the best performance in identifying strong (74.3-94.7% occupancy), median (53.8-74.3%), weak (33.3-53.8%), and non-binding (12.8-33.3%) sites. Chol-GAT ranked second, followed by Chol-GCN, whereas Chol-3DCNN model showed no correlation with MD-simulated occupancy. We speculate that, because the transmembrane domain of the PIEZO2 channel is substantially larger than those of the proteins curated from the PDB database, the binding sites are likely to be more complex and diverse. Chol-GAT model may benefit more from the complexity and thus performed better for PIEZO2 than it did in the positive test set or the external validation set.

**Figure 9.**
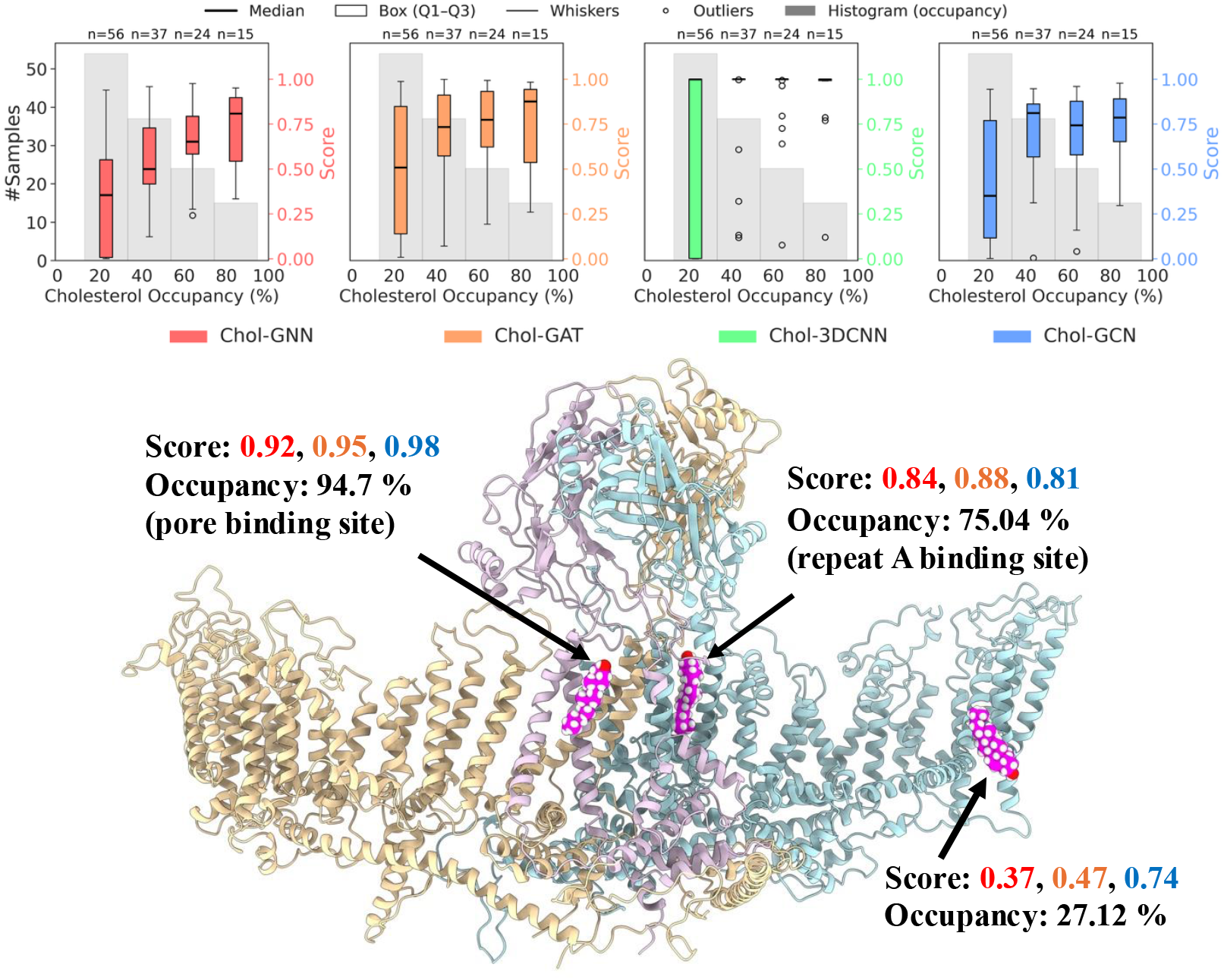
Comparison of predicted cholesterol-binding scores from Chol-GNN, Chol-GAT, Chol-3DCNN, Chol-GCN with MD-simulated cholesterol occupancy for PIEZO2. Box plots show the distribution of model prediction scores across four occupancy bins (Bin 1: 12.8–33.3%, n=56; Bin 2: 33.3–53.8%, n=37; Bin 3: 53.8–74.3%, n=24; Bin 4: 74.3–94.7%, n=15), averaged over two MD simulation replicas, with gray histograms indicating the number of samples in each bin. Representative PIEZO2 snapshots highlight three cholesterol-binding sites (magenta) with high, intermediate, and low occupancies from MD simulations. Corresponding model prediction scores (Chol-GNN, Chol-GAT, Chol-GCN) and occupancies are indicated for each site.

Among the strong cholesterol binding sites predicted by both our all-atom MD simulations and CholBindNet, one site is located in the PIEZO2 repeat A region. This site was previously reported in a coarse-grained Martini simulation study of PIEZO2 in the closed conformation [26]. For this site, our model prediction scores are 0.84 for Chol-GNN, 0.88 for Chol-GAT, and 0.81 for Chol-GCN, and MD-simulated occupancy of 75.04% (**Figure 9** bottom). In the absence of an experimentally determined binding site, the concordance among deep-learning predictions, all-atom MD simulations, and coarse-grained MD simulations provides strong confidence in our model’s prediction. In addition, our all-atom MD simulations of the PIEZO2 open state here also reveal a new cholesterol binding site near the pore, whose functional relevance has yet to be experimentally validated.

Although the qualitative agreement is encouraging, the deep-learning model and physics-based MD simulations are not expected to agree entirely. The deep-learning model relies on single snapshots from MD trajectories and therefore does not capture the full dynamics of the binding pockets. Additionally, the model focuses solely on the local binding environment and does not account for the binding pathway. Nevertheless, the qualitative agreement between our deep-learning models and MD simulations suggests that our model benefits from the PU-learning algorithm, enabling it to differentiate between strong and weak binding sites rather than functioning merely as a binary classifier.

## 3 Discussion

Cholesterol is emerging as a central regulator in the pathological mechanisms underlying cancer, neurodegeneration, and stroke. Predicting cholesterol binding sites remains challenging due to its highly “undrug-like” physicochemical properties, making cholesterol interactions difficult to probe experimentally and computationally. However, recent advances in cryo-EM structure determination of membrane proteins, combined with the rapid development of deep-learning pattern-recognition algorithms, now offer an unprecedented opportunity to study and predict cholesterol binding with greater accuracy. The classical CRAC and CARC motifs, long been used to describe cholesterol-binding sites, have failed to capture the local 3D environment. Our atom-based model bypasses these limitations by learning directly from atomic coordinates and subtypes, without imposing motif assumptions or introducing bias from predefined binding-site classifications. The dataset used in this study, comprising more than 800 high-resolution cholesterol-binding structures, represents, to our knowledge, the largest cholesterol-specific training set assembled to date.

A unique strength of our approach lies in its semi-supervised training on unlabeled data generated by AutoDock Vina. Because Vina was not parameterized for cholesterol and thus produces many false-positive sites, incorporating both experimentally validated cryo-EM binding sites and Vina-generated candidate sites enables the model to learn subtle distinctions between true cholesterol-binding patterns and generic hydrophobic pockets. This hybrid training strategy enables graphic models to recognize cholesterol-specific binding environments beyond what can be captured from sequence motifs or standard docking scores alone.

Our model significantly outperforms three state-of-the-art, generic deep-learning ligand-binding prediction models on the cholesterol dataset, underscoring the need for task-specific models that are explicitly designed and trained for cholesterol binding. That said, we can not rule out the possibility that generic ligand-binding models could improve their performance through fine-tuning on our curated dataset. Importantly, however, the ranking power among strong to weak binding sites, derived from our PU-learning framework, and demonstrated here using the PIEZO2 MD simulation data, offers a unique advantage, particularly because cholesterol binding is more promiscuous and structurally diverse in large transmembrane proteins.

Another advantage of our approach is the model’s interpretability. The feature importance, derived solely from atomic distances and atom subtypes, consistently highlights branched apolar and aromatic residues, echoing the key elements of the CRAC/CARC motifs while providing atom-level specificity. This level of detail allows us to understand why the model classifies a site as a strong cholesterol-binding site and which residue pairs and spatial distances contribute most to the binding. The divergence observed in the GAT model reflects its tendency to assign greater weights to non-local interactions, an ability that becomes useful when binding sites are more complex and heterogeneous, such as in PIEZO2. Together, these insights and interpretability open new avenues for integrating cholesterol into the therapeutic landscape, revealing disease-related mechanisms and enabling the identification of cholesterol-binding sites that can be leveraged for the rational design of novel drug modalities.

Although our models outperform existing deep-learning approaches for predicting cholesterol binding sites, this task is far from complete. The simplicity of our current feature representation leaves room for further improvement. For example, our model uses only protein atom features. While this is sufficient to define a binding pocket, it loses information about the ligand’s binding pose. Unlike generic ligands, cholesterol is an integral component of the lipid bilayer and adopts opposite orientations in the outer and inner leaflets. We observed that Vina and DiffDock occasionally place cholesterol in the correct binding region but in the wrong orientation, because there is no membrane environment present to constrain the orientation of the sterol. Incorporating cholesterol atom coordinates and atom-type features directly into the node representation could allow the model to capture pose-specific information and further enhance predictive accuracy. Additionally, a distance cutoff of 5 Å is a commonly used criterion to define protein atoms that directly bind ligands. In future work, multiple distance cutoffs could be incorporated to capture longer-range protein–cholesterol interactions.

Our dataset of ∼800 experimental cholesterol-bound protein structures remains relatively small and consists only of true positives. In the past decade, MD studies of membrane proteins have increasingly shifted from pure POPC bilayers to multicomponent membranes that frequently include cholesterol. Harnessing dynamically sampled cholesterol binding events from MD trajectories will enable the development of models with quantitative ranking power approaching that of computationally expensive MD simulations, providing a richer and more nuanced description of cholesterol–protein interactions.

## 4 Methods

### 4.1 Model Architecture

The *CholBindNet* framework integrates four complementary deep learning architectures: (1) CholBind Graph Neural Network (Chol-GNN), (2) CholBind Graph Convolutional Network (Chol-GCN), (3) CholBind Graph Attention Network (Chol-GAT), and (4) CholBind Three-Dimensional Convolutional Neural Network (Chol-3DCNN). For the Chol-GNN, Chol-GCN, and Chol-GAT models, the adjacency matrix is constructed using structural information derived from the inverse distance matrix, while atom subtypes (such as CA, CB, N, O, and S) are encoded as node features (**Table 1**). In the Chol-3DCNN model, each atom is represented as a voxel within a spatial grid defined by the protein’s structural coordinates, enabling volumetric feature extraction. Detailed architecture is illustrated in **Figure 10**, and each component of network architecture and its implementation are provided in the following sections.

**Figure 10.**
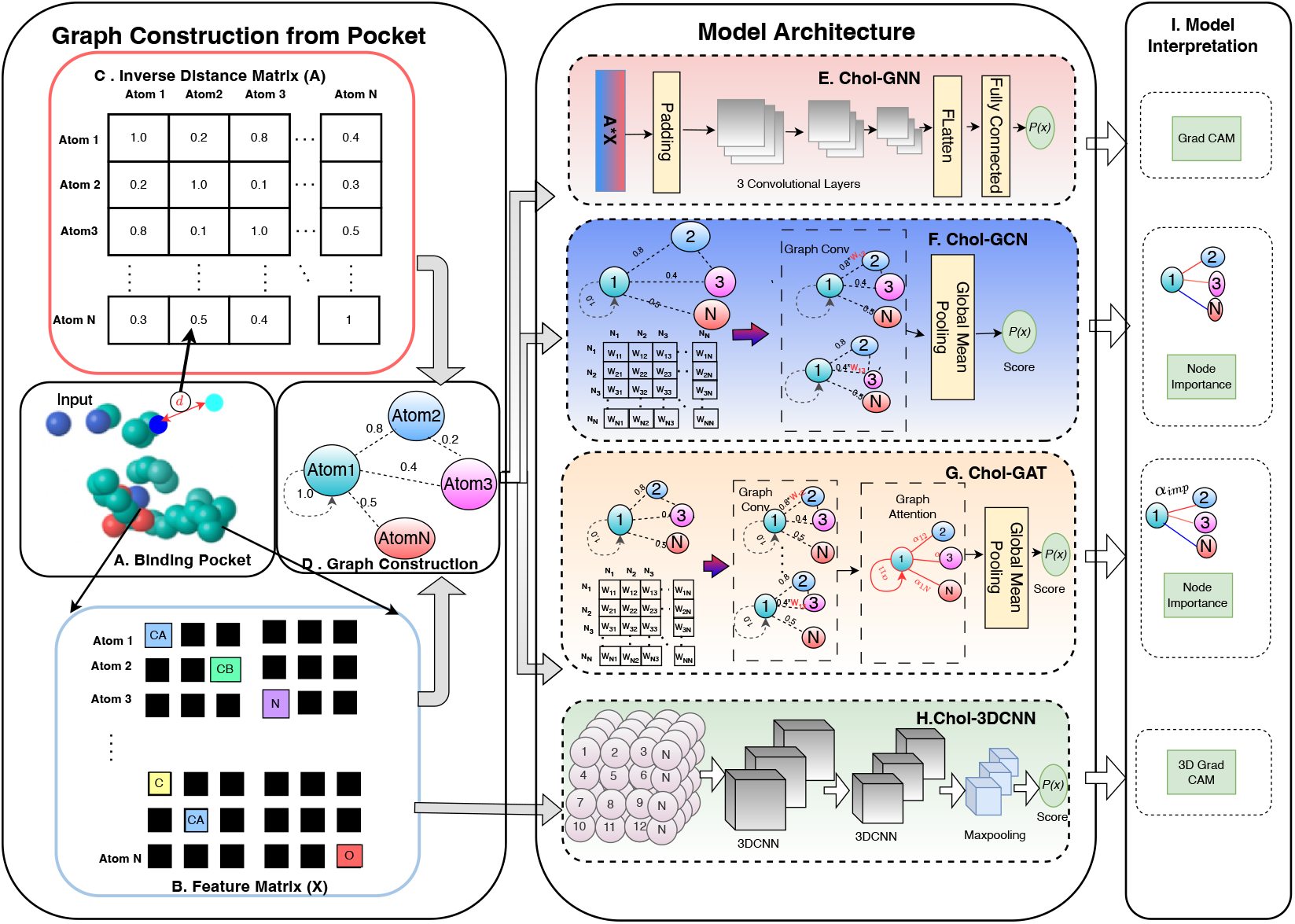
Detailed overview of Chol-Bind Framework; (A) Binding pocket (Filtered PDB;input); (B) Inverse Distance Matrix; (C) Feature Matrix (One-hot encoding); (D) Graph Construction; (E) Chol-Graph Neural Network architecture; (F) Chol-Graph Convolutional Neural Networks; (G) Chol-Bind Graph Attention Neural Network; (H) Chol-3D Convolutional Neural Network; and (I) Model Interpretation; Chol-GNN and Chol-3DCNN gives atom-subtype importance, and Chol-GCN and Chol-GAT gives us residue importance along side of atom subtype importance (see methods)

#### Inverse-distance matrix

To capture the structural information of the protein, inter-atomic relationships are represented using an inverse-distance matrix. Here, an inter-atomic relationship refers to the spatial and geometric dependency between pairs of atoms that collectively define the local and global three-dimensional organization of the protein structure. Such relationships are essential for modeling how atomic arrangements influence molecular interactions and potential binding events.

Given the three-dimensional Cartesian coordinates of each atom, denoted by **r**_*i*_ ∈ ℝ^3^ for the *i*^th^ atom, the pairwise Euclidean distance between atoms *i* and *j* is defined as

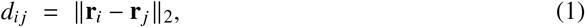

where ∥·∥_2_ denotes the Euclidean (L2) norm, and *d*_*ij*_ measures the spatial separation between atoms *i* and *j* in the protein’s three-dimensional structure.

Based on these distances, the adjacency matrix (inverse-distance affinity) **A** ∈ ℝ^*N*×*N*^ is constructed, where *N* is the total number of atoms. Each element *A*_*ij*_ encodes the structural proximity between atoms *i* and *j* as

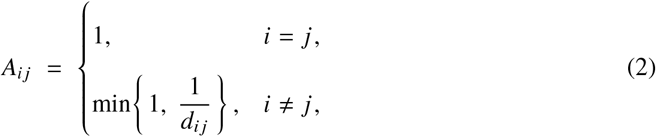

such that atoms in close spatial proximity exhibit higher affinity values, while distant atoms contribute weaker connections. The min operation caps the maximum affinity at 1, ensuring numerical stability and preventing extremely small inter-atomic distances from dominating the representation. This inverse-distance formulation quantitatively encodes the geometric topology of the protein and provides a continuous, structure-aware adjacency representation suitable for graph-based neural architectures (Chol-GNN, Chol-GCN, and Chol-GAT) to learn spatial dependencies and binding-relevant structural patterns. The resulting affinity matrix **A** serves as the foundation for graph construction, defining connectivity among atoms and guiding feature propagation in subsequent deep learning modules.

### 4.2 Graph construction

Using the adjacency matrix **A**, a protein is represented as a graph **G** = (**A, X**) to facilitate structural learning through graph-based neural networks. In this representation, each node corresponds to an atom, and edges represent structural connections weighted by spatial proximity as defined in **A**. The node feature matrix **X** ∈ ℝ^*N*×*F*^ encodes atom-level physicochemical information, where *N* denotes the number of atoms and *F* represents the number of feature channels. Each atom is described using a one-hot encoding of its subtype (e.g., CA, CB, N, O, S), which may be optionally augmented with additional features such as atomic charge, hybridization state, or residue-level identifiers (**Table S1**).

This formulation allows the graph representation to capture both topological and geometric information inherent to the protein structure. The adjacency matrix **A** embeds structural context by encoding pairwise spatial dependencies, while the feature matrix **X** captures local chemical identity. Together, these components provide a comprehensive, structure-aware input representation that enables the Chol-GNN, Chol-GCN, and Chol-GAT architectures in CholBindNet to learn spatially informed embeddings of atomic environments relevant to cholesterol-binding site identification.

#### 4.2.1 CholBind Graph Neural Network (Chol-GNN)

The graph network module in Chol-GNN integrates geometric and chemical features of proteins to identify cholesterol-binding sites. From the filtered set of atoms, two matrices are constructed: the inverse-distance matrix **A** and the atom feature matrix **X**. The matrix **A** ∈ ℝ^*N*×*N*^ encodes spatial relationships among all *N* atoms in the protein, where each entry *A*_*ij*_ represents the inverse of the Euclidean distance between atoms *i* and *j*, with *A*_*ii*_ = 1 denoting self-connections. The matrix **X** ∈ ℝ^*N*×*F*^ encodes *F* atom-level features for each of the *N* atoms, including element type and other physicochemical attributes, represented using one-hot encoding.

To generate a structured representation suitable for convolutional processing, an interaction matrix **M** is computed as

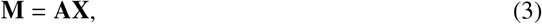

where **M** ∈ ℝ^*N*×*F*^ aggregates atom features weighted by their spatial affinities. Each element *M*_*ij*_ thus reflects how the *j* ^th^ feature of atom *i* is influenced by neighboring atoms in proportion to their geometric proximity, effectively embedding the protein’s three-dimensional topology into a two-dimensional tensor.

This matrix **M** is treated as an image-like input to a convolutional backbone designed to extract hierarchical spatial features. The network comprises multiple convolutional layers with batch normalization, nonlinear activation, and max-pooling, which progressively learn higher-order structural patterns. The resulting feature maps are flattened and passed through fully connected layers with dropout regularization, producing a final scalar logit (converted to a probability via a sigmoid during training/evaluation) that reflects the predicted likelihood of cholesterol binding. This architecture enables CholBindNet to jointly model local atomic interactions and global structural organization while preserving interpretability through spatially weighted feature aggregation.

#### 4.2.2 CholBind Graph-Convolutional (Chol-GCN)

The CholBind Graph-Convolutional Network (Chol-GCN) module in CholBindNet takes inverse-distance adjacency matrix **A** and the atom feature matrix **X** as input similar to Chol-GNN. The matrix **A** ∈ ℝ^*N*×*N*^ is interpreted as a weighted adjacency, where larger values correspond to shorter inter-atomic distances, and thus stronger spatial affinities. The matrix **X** ∈ ℝ^*N*×*F*^ encodes *F* atom-level features for each of the *N* atoms, representing their chemical identities and local physicochemical descriptors.

Each GCN layer propagates information across the molecular graph by aggregating messages from spatially neighboring atoms. The degree-normalized propagation rule for the *l*^th^ layer is given by

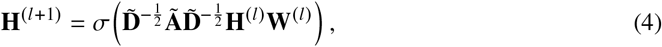

where **Ã** = **A** + **I**_*N*_ is the adjacency matrix with self-loops added. Self-loops are automatically incorporated by the GCNConv operator to preserve node-specific information during message passing. 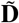 is the corresponding degree matrix with entries 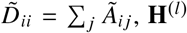 is the input node embedding at layer *l*, **W**^(*l*)^ is a learnable weight matrix, and *σ*(·) denotes a nonlinear activation function. This normalization ensures numerical stability and prevents nodes with many connections from dominating the aggregation. The inverse-distance weights in **A** modulate message strength, emphasizing contributions from atoms that are spatially closer.

Successive GCN layers apply this propagation to learn higher-order spatial representations of the molecular structure. The resulting node embeddings are pooled into a graph-level representation using global mean pooling, followed by fully connected layers that integrate global contextual information. The network outputs a single scalar logit, which is transformed into a probability via a sigmoid function during training and evaluation. This formulation enables CholBindNet to capture both the geometric proximity and the chemical environment of atoms, providing a physically grounded representation of cholesterol–protein interactions without the need for explicit attention mechanisms.

#### 4.2.3 CholBind Graph-Attention (Chol-GAT)

The CholBindNet Graph Attention Network module (Chol-GAT) extends the graph convolutional paradigm by incorporating a learnable attention mechanism to adaptively weight atomic interactions. As in the Chol-GNN and Chol-GCN modules, the inputs consist of the inverse-distance adjacency matrix **A** and the atom feature matrix **X**. The matrix **A** ∈ ℝ^*N*×*N*^ encodes spatial proximity between atoms, while **X** ∈ ℝ^*N*×*F*^ represents *F* atom-level features such as element type and local chemical descriptors.

In contrast to fixed, degree-normalized aggregation in standard GCNs, Chol-GAT model learns an attention coefficient *α*_*ij*_ for each edge (*i, j*) that determines the relative contribution of atom *j* to the updated representation of atom *i*. For a given layer *l*, the propagation rule is expressed as

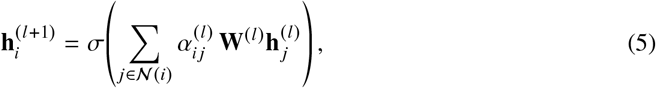

where 𝒩 (*i*) denotes the set of neighbors of atom *i*, **W**^(*l*)^ is a learnable weight matrix, and *σ*(·) is a nonlinear activation function. The attention coefficients 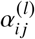 are computed using a shared attention mechanism,

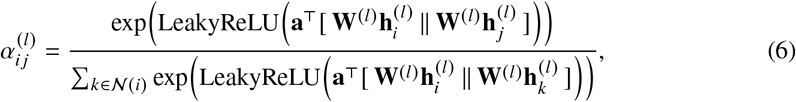

where **a** is a learnable attention vector and ∥ denotes vector concatenation. This formulation allows the model to dynamically focus on the most chemically or spatially relevant neighboring atoms, guided implicitly by both learned feature representations and the structural weights in **A**.

The node embeddings from the final attention layer are aggregated through global mean pooling to generate a graph-level representation. This embedding is subsequently processed through batch normalization, dropout regularization, and a fully connected output layer with sigmoid activation to yield the probability of cholesterol-binding. By assigning adaptive importance to atom–atom interactions, the GAT module enhances CholBindNet’s capacity to identify critical substructures and interpret the molecular determinants underlying cholesterol–protein binding.

### 4.3 Chol-Bind 3D Convolutional Neural Network (Chol-3DCNN)

In the Chol-Bind three-dimensional convolutional neural network (Chol-3DCNN) module, each atom is first encoded according to its subtype and then mapped to a discrete three-dimensional grid based on its Cartesian coordinates. The molecular volume is represented using a 30 × 30 × 30 voxel grid, which was selected after initial experiments with a 20^3^ grid revealed overlapping atom positions. When two atoms were located within 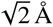 of each other, one atom was shifted to the nearest unoccupied voxel to preserve spatial uniqueness. To increase data diversity, each structure was randomly rotated five times, and the rotated versions were used as independent training samples. This rotation-based augmentation is a key advantage of the 3D-CNN model, as it allows the network to learn spatial invariance without relying on predefined graph connectivity.

The input tensor **V** ∈ ℝ^*C*×30×30×30^ represents the voxelized molecular structure, where *C* = 37 corresponds to the number of atom subtypes used as input channels. The 3D-CNN applies a sequence of volumetric convolutional layers to extract hierarchical spatial features. The first convolutional block uses 1 × 1 × 1 kernels with 64 filters, followed by a ReLU activation and 2 × 2 × 2 max pooling to downsample the feature map. Subsequent blocks employ 3 × 3 × 3 kernels with 64 and 128 filters, each followed by ReLU activation and max pooling operations to progressively capture local geometric patterns and global spatial organization.

The resulting 3D feature map is flattened and passed through two fully connected layers with 256 and 128 hidden units, each followed by ReLU activation, culminating in a final fully connected layer with two output neurons and a softmax activation function. The model outputs a categorical probability distribution indicating whether the structure represents a cholesterol-binding site or a non-binding region. This volumetric formulation enables CholBindNet to directly learn from the three-dimensional spatial arrangement of atoms, capturing complex geometric dependencies that are not explicitly modeled in the graph-based architectures.

### 4.4 Positive-Unlabedling learning objective

CholBindNet is trained within a positive–unlabeled learning framework rather than a conventional binary classification setting, because true negative cholesterol-binding sites cannot be reliably identified [28]. To address this challenge, the PU strategy constructs an effective binary decision model using only confirmed positives and unlabeled samples. This is accomplished through a binning procedure in which multiple weakly supervised classifiers are trained on distinct sampled subsets of the data and later combined into a single consensus model, as illustrated in (**Figure 1b**). After aggregation, a percentile-based labeling scheme is applied to assign each predicted pocket to one of four categories: positive, pseudo-positive, pseudo-negative, or negative.

In this PU framework, a small subset of confirmed positive samples is deliberately masked by relabeling them as unlabeled and inserting them into the larger unlabeled pool. These masked positives, referred to as spies, provide an internal reference for assessing model behavior. The key idea is that if the model can reliably recover spies as positives despite their unlabeled status, then the PU strategy is effective at separating potential binding pockets from likely nonbinders within the unlabeled data.

The unlabeled pool therefore consists of a heterogeneous collection of binding pockets that may contain true but unknown cholesterol-binding sites, nonbinding sites, and the inserted spy binding sites samples. To construct the binning ensemble, the unlabeled set *D*_*u*_ is randomly partitioned into *M* subsets, 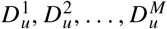, where each subset 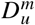 serves as the unlabeled component for the *m*-th bin. For each bin, the training set *D*^*m*^ is created by combining the positive training set 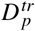 (True-positive binding after removing spies) with the corresponding unlabeled subset:

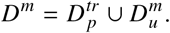

A separate classifier is trained on each *D*^*m*^, treating the unlabeled subset 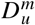 as negative for that bin. After training all *M* classifiers, the average posterior probability for a sample *x* is computed as

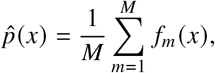

where *f*_*m*_ (*x*) denotes the posterior probability assigned to sample *x* by the classifier trained on the *m*-th bin. The aggregated score 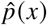 provides a consensus estimate of the likelihood that *x* is a cholesterol-binding pocket and is used for the final percentile-based labeling procedure. To convert the aggregated ensemble scores into adaptive supervision labels, we implemented a percentile-based procedure driven by the empirical distribution of spy scores. Let 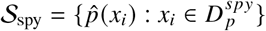 denote the aggregated probability scores assigned to all spy samples. The quartiles *P*_25_, *P*_50_, and *P*_75_ are computed from 𝒮_spy_ and used as decision boundaries for labeling all samples in the unlabeled pool. The full procedure is outlined below.

#### Algorithm 1

Percentile-based Labeling Using Spy Score Distribution

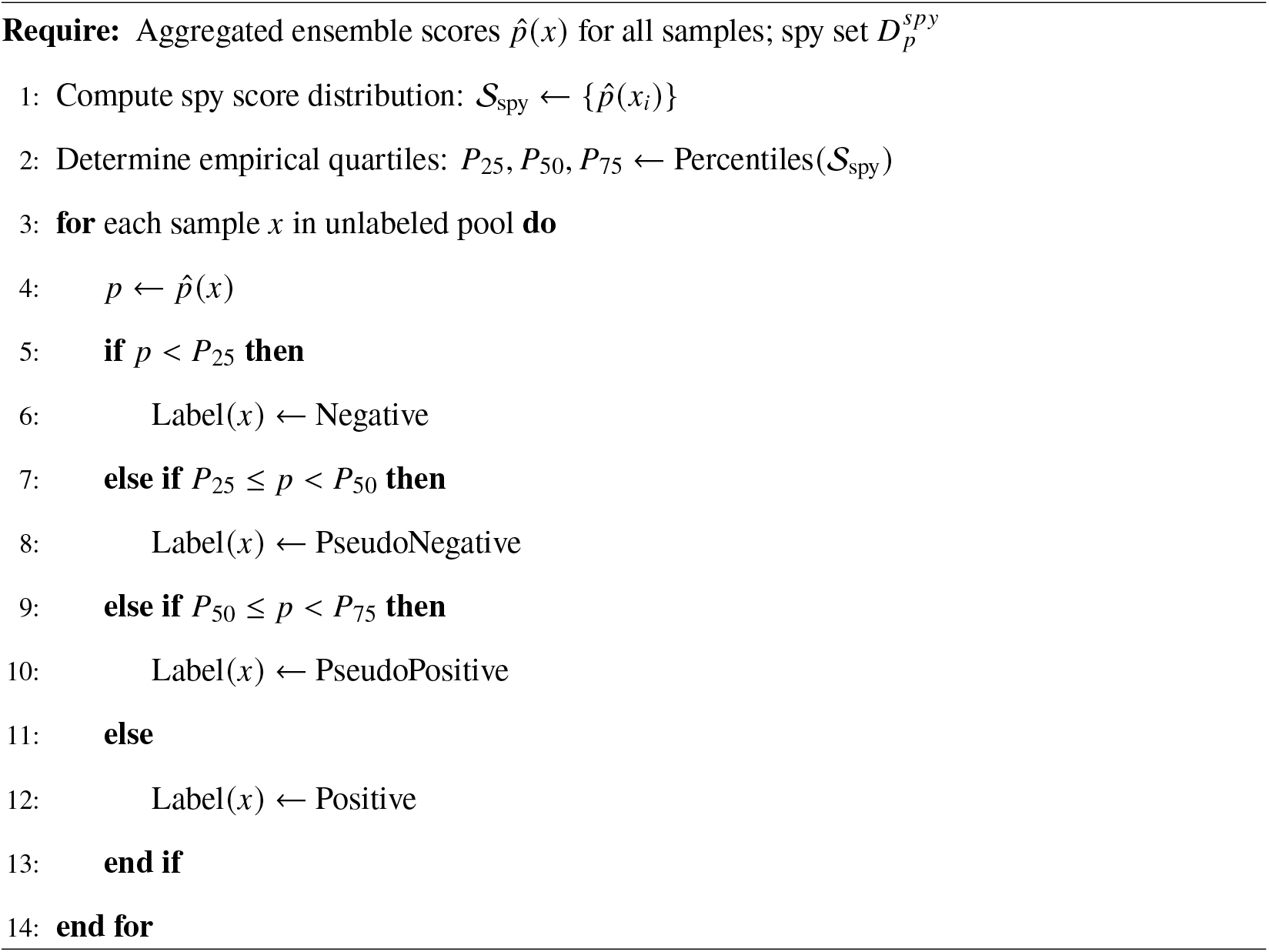

This procedure generates adaptive, distribution-aware labels that characterize each sample according to its relative position within the spy-derived score distribution. Because the thresholds are defined by the empirical quartiles of 𝒮_spy_, the labeling remains robust to calibration differences across individual bin classifiers and adjusts naturally to variations in dataset difficulty.

### 4.5 Model Interpretation

#### 4.5.1 Interpretability of the Chol-GNN module

To identify which atom subtypes contribute most strongly to the cholesterol-binding prediction, saliency analysis was applied to Chol-GNN. Recall that the interaction matrix **M** = **AX** combines spatial and chemical information by weighting atom features **X** with the inverse-distance affinities encoded in **A**. Each row **M**_*i*_ therefore represents the effective feature vector of atom *i* after aggregating information from its neighboring atoms. Because Chol-GNN employs a convolutional backbone operating on **M** rather than on an explicit edge list, interpretability is limited to atom-level or feature-channel attributions rather than specific atom–atom interactions.

The saliency of each atom was quantified using the gradient-based importance measure

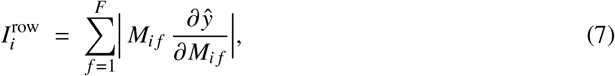

where *M*_*if*_ denotes the contribution of the *f* ^th^ feature of atom *i* in the interaction matrix **M**, *F* is the number of features per atom, and *ŷ* is the model’s predicted probability of cholesterol binding. The partial derivative 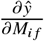 measures the sensitivity of the prediction to perturbations in that specific feature, while the product 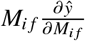 captures both feature magnitude and its influence on the output. Summing the absolute values across all feature dimensions yields the overall saliency score 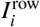, reflecting the importance of atom *i* to the model’s decision.

Atoms with higher 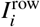 values are interpreted as key contributors to the cholesterol-binding prediction, highlighting atom subtypes or regions that play dominant structural or chemical roles. This formulation enables localized interpretation of the Chol-GNN’s decision process while preserving the structural abstraction inherent to the convolutional formulation.

#### 4.5.2 Interpretability of the Chol-GCN module

In Chol-GCN, information is propagated across atoms using fixed inverse-distance weights *A*_*ij*_ combined with learned linear filters. Because these weights directly encode spatial proximity, interpretability can be derived from both edge-level and node-level sensitivities, allowing identification of influential atom pairs and individual residues involved in cholesterol binding.

Edge-level importance is quantified using a gradient-based saliency score defined as

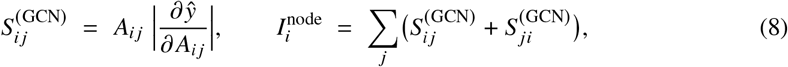

where *ŷ* denotes the model’s predicted probability of cholesterol binding, *A*_*ij*_ is the inverse-distance affinity between atoms *i* and *j*, and **x**_*i*_ represents the feature vector of atom *i*. The derivative 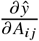 measures how sensitive the model’s output is to small perturbations in the edge weight *A*_*ij*_, thereby capturing the influence of the spatial relationship between atoms *i* and *j* on the prediction. The product 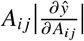 combines geometric proximity and predictive sensitivity to yield the edge-level importance score 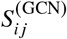.

Node-level importance can be computed by aggregating the saliency contributions of all edges incident to node *i*, through 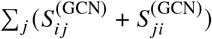. Atoms with higher 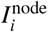 values are interpreted as having a greater impact on the final prediction. Because *A*_*ij*_ explicitly encodes physical proximity, these explanations naturally align with molecular geometry, which indicates that atoms spatially closer to the binding pocket contribute more strongly, while still distinguishing which specific neighboring interactions were most influential in determining the model’s output.

#### 4.5.3 Interpretability of the Chol-GAT module

Chol-GAT provides fine-grained, interaction-level interpretability by assigning learned attention weights to edges between atoms. Each directed edge (*i* → *j*) in the molecular graph is associated with an attention coefficient *α*_*ij*_, which reflects the relative importance of atom *j* ‘s features to the representation update of atom *i*. These coefficients are learned during training and are returned alongside the edge index, enabling direct mapping of model attention to specific atom–atom interactions.

To quantify the contribution of each interaction to the final prediction, an attention-based gradient score, denoted as *Attention*×*Gradient*, is computed as

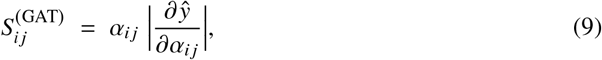

where *ŷ* represents the model’s predicted probability of cholesterol binding, and 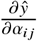 is the gradient of the output with respect to the attention coefficient. The magnitude of this term captures the sensitivity of the prediction to changes in *α*_*ij*_, while scaling by *α*_*ij*_ incorporates the intrinsic attention assigned to that edge. The resulting 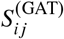 score therefore highlights atom pairs that are both strongly attended to and influential in determining the model’s output.

To compute atom-level importance, edge saliency values are aggregated across all edges incident to each atom:

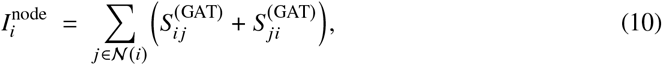

where 𝒩 (*i*) denotes the set of neighboring atoms connected to atom *i*. The resulting 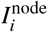 quantifies each atom’s cumulative influence through its interactions. Higher values indicate atoms that participate in multiple critical or strongly weighted connections within the binding pocket.

Although attention values *α*_*ij*_ are not causal proofs of importance, they provide interpretable hypotheses regarding key interactions. To corroborate these findings, targeted edge ablation experiments are performed by removing the top-*k* edges ranked by 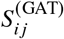 and measuring the corresponding decrease in *ŷ*. This validation step helps distinguish genuinely influential atom pairs from spurious correlations, enhancing interpretability and trustworthiness in CholBindNet’s predictions.

#### 4.5.4 Interpretability of the Chol-3DCNN module

Chol-3DCNN operates on a voxelized representation of the protein structure with 37 input channels, each corresponding to a specific atom subtype. Interpretability is achieved through a three-dimensional extension of Gradient-weighted Class Activation Mapping (Grad-CAM), which identifies the spatial regions within the 3D grid that most strongly influence the model’s output.

Let 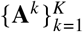 denote the set of 3D activation maps from the final convolutional layer, where **A**^*k*^ ∈ ℝ^*D*^′×*H*′×*W*′ represents the activation volume for the *k*^th^ feature channel, and *K* is the number of such channels. The scalar output of the network, *ŷ*, corresponds to the logit or post-sigmoid probability of cholesterol binding. To compute Grad-CAM weights, the gradient of *ŷ* with respect to each activation map is averaged spatially:

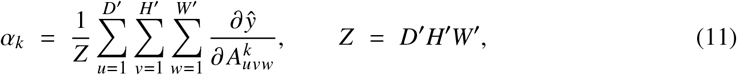

where *α*_*k*_ measures the importance of feature channel *k*, and *Z* is the total number of spatial locations. The resulting class-activation map is obtained as

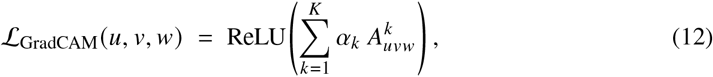

where ReLU ensures that only features positively contributing to the target class are retained. The resulting 3D saliency volume ℒ_GradCAM_ highlights spatial regions that most strongly support a positive binding prediction.

For interpretability and visualization, ℒ_GradCAM_ is normalized to the range [0, 1] as

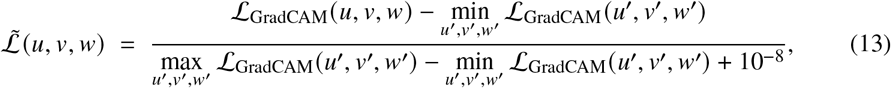

where the small constant ensures numerical stability. The normalized map is then upsampled to match the input resolution, allowing voxel-level attributions to be aggregated to the atom level. For each atom *a*, occupying a set of voxels *V* (*a*), an atom-level importance score is computed as

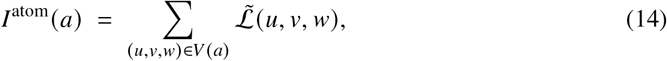

which quantifies the cumulative saliency over all voxels associated with that atom.

This approach produces spatially localized explanations that highlight the 3D regions—and, consequently, the corresponding atoms—that most strongly support a positive classification. Unlike graph-based models such as GAT or GCN, which explicitly model pairwise atomic interactions, the 3D-CNN provides voxel- and atom-level importance maps that emphasize geometric regions rather than specific edges. This makes it particularly effective for identifying volumetric structural motifs relevant to cholesterol-binding phenomena.

#### 4.5.5 Ranking of Interpretable Structural Features

To systematically rank the most important structural features identified by the interpretability analyses, we applied a unified ranking procedure across the Chol-GCN, Chol-GAT, Chol-GNN, and Chol-3DCNN models. For the graph-based models (Chol-GCN and Chol-GAT), edge-level importance scores were computed using the corresponding gradient-based interpretation formulations. Within each sample, edge importance scores were normalized such that the highest-scoring edge was assigned a value of 1, ensuring comparability across samples and mitigating scale differences between predictions.

Although interpretation is computed at the atom–atom edge level, ranking is ultimately performed at the residue–distance level to enable biologically meaningful analysis. For each important atom–atom edge, the corresponding residue identities were retrieved, and the inter-residue distance was assigned to one of five distance bins: 4–8 Å, 8–12 Å, 12–16 Å, 16–20 Å, and >20 Å. Edges connecting atoms within the same residue were ignored to focus exclusively on unique residue–residue interactions. Importance scores were then aggregated across samples to rank the most influential residue–distance pairs.

To rank individual residues and atom subtypes, a frequency-based approach was used across all four models. For each sample, the top five most important features were identified—corresponding to edges for Chol-GCN and Chol-GAT, grid points for Chol-3DCNN, and rows of the interaction matrix for Chol-GNN. The frequency with which each residue or atom subtype appeared among these top-ranked features was then counted across all samples, and residues or subtypes were ranked accordingly. This approach highlights consistently influential structural elements across models while reducing sensitivity to single-sample outliers.

### 4.6 Molecular docking protocol

The molecular docking workflow was implemented using custom Python scripts incorporating the BioPython library [29] for structural manipulation, along with external utilities for file format conversion and postprocessing. The pipeline consisted of four main stages: (i) protein structure preparation, (ii) grid box determination, (iii) receptor and ligand preparation, and (iv) post-docking analysis.

#### 4.6.1 Protein Structure Preparation and Cleaning

High resolution protein structures (resolution < 3.5 Å) was obtained from the Protein Data Bank (PDB). The structure, originally provided in CIF format, was converted to PDB format using BioPython [29]. Nonstandard residues, heteroatoms, and cholesterol molecules were removed to produce a clean receptor model suitable for docking studies.

#### 4.6.2 Docking grid box determination

For blind docking using AutoDock Vina [30], the docking grid was programmatically defined to encompass the entire receptor. The grid boundaries were calculated from the minimum and maximum atomic coordinates along the x, y, and z axes. The grid center was set at the geometric midpoint, and a uniform padding of 10 Å was applied along each axis to ensure complete coverage of the receptor surface.

#### 4.6.3 Receptor and Ligand Preparation

The processed receptor structures were converted to PDBQT format, the required input for AutoDock Vina [30], using AutoDockFR 1.0 (ADFR) [31]. The cholesterol ligand was retrieved from PubChem in SDF format and converted to PDBQT using Meeko [32].

#### 4.6.4 Molecular Docking

Docking simulations were carried out using AutoDock Vina (v1.2.3) in a high-throughput mode. A custom Python script automated the docking process across all receptor structures. For each receptor, a configuration file was generated dynamically, containing grid box parameters and relevant docking settings derived from the previous stages.

#### 4.6.5 Post-Docking Analysis

Following docking, output files were analyzed to extract 4 to 5 distinct top-ranked docking poses. The docked complexes were converted into standard chemical formats using Meeko for visualization and further structural analysis [32].

### 4.7 Benchmarking Deep learning Methods

#### 4.7.1 LABind Preparation

LABind v1.0.0-alpha was evaluated for its performance on the external validation dataset. Cholesterol SMILES strings and individual protein chains from the external validation dataset were extracted. Experimental PDB structures with its FASTA sequences were employed directly without using ESMFold. For each protein, only chains containing residues located within 5 Å of cholesterol in the experimental complex were considered relevant for binding. Because LABind accepts only a single chain as input, we excluded cholesterol binding sites formed by more than one chain. Thus, a subset of five structures, each containing exactly one cholesterol binding chain, was used for evaluation. LABind’s default probability threshold of 0.48 was applied to classify residues as high probability binding site residues. These residues were then compared with experimentally observed binding residues, defined as those within 5 Å of cholesterol. If at least one of these predicted residues matched any of the experimentally observed residues, we considered it as positive.

#### 4.7.2 Chai-1 Preparation

Chai-1 v0.6.1 was also evaluated for its performance on the external validation dataset. Required inputs are the FASTA sequences of all protein chains contributing to cholestering binding along with the SMILES string of cholesterol. Unlike LABind, Chai-1 accepts multichain protein inputs. However limited GPU memory restricts us to use only a subset of 23 structures with residue numbers less than 1000. The model outputs a predicted ligand pose. To maintain a binary classification consistent with LABind, predictions were evaluated by determining whether the center of mass (COM) of the predicted cholesterol pose is within 5 Å of the COM of the experimenntal pose.

#### 4.7.3 DiffDock Preparation

DiffDock v1.1.3 was evaluated using full protein structures in the external validation dataset. Similar to Chai-1, DiffDock supports multichain inputs. Because DiffDock does not perform co-folding, no hardware limitation, hence all structures in the external validation dataset were evaluated. DiffDock outputs predicted cholesterol poses, which were assessed using the same COM distance metric applied to Chai-1.

### 4.8 All-atom molecular dynamics of PIEZO2 channel

The PIEZO2 model used in this study corresponds to a fully open state conformation generated from the cryo-EM structure of mouse PIEZO2 (PDB ID: 6KG7) [33] using our hybrid molecular dynamics simulations [34]. To reduce the computational cost, the peripheral portion of three arms of PIEZO2 protein were truncated by deleting the THU1–THU4 helical bundles. The systems were assembled using the CHARMM-GUI Membrane Builder web server [35]. The CHARMM36m force field was applied to the protein, while the CHARMM36 force field was used for the lipids [36]. The percent composition of the inner and outer leaflets was POPC (65/70), cholesterol (20/20), phosphatidic acid (10/10), and PIP2 (5/0). Potassium ions were added to neutralize the system, and TIP3P water model was used to solvate the system. The total system contained 1,638,978 atoms, with box dimensions of 318.09 × 318.09 × 170 Å^3^.

The equilibrium all-atom MD simulations were performed using GROMACS version 2024.2-smp [37]. Energy minimization was performed using 10,000 steps of the steepest descent algorithm. Non-bonded interactions were computed with long-range electrostatics handled by the particle mesh Ewald (PME) method [38], while van der Waals interactions were truncated at 1.2 nm using a force-switch modifier applied between 1.0–1.2 nm to smoothly approach the cutoff . All bond lengths involving hydrogen atoms were constrained by using the LINCS algorithm [39], allowing for a 2 fs timestep in the subsequent simulations. During equilibration at 310.15 K, positional restraints were applied to the protein backbone, side chains, and lipids, along with dihedral restraints. The Nose-Hoover thermostat was used for temperature coupling, with separate coupling groups for the protein, membrane, and solvent (*τ* = 1.0 ps) [40]. Semi-isotropic pressure coupling was applied in the later equilibration stages using the Parrinello-Rahman barostat at 1 bar, with a compressibility of 4.5 × 10^−5^ bar^−1^ and a relaxation constant of 5.0 ps [41, 42].

For production simulation, the final gromacs gro coordinate file was uploaded to the ANTON3 supercomputer and converted into the Desmond chemical format using Viparr [43]. Simulations were performed using a multigrator integrator scheme with a timestep of 0.002 ps. Temperature control was achieved with a Nose–Hoover thermostat, and pressure was regulated using the Martyna–Tuckerman–Klein (MTK) barostat [40, 44], with a target temperature of 310.15 K and a relaxation time of 0.0417 ps. Semi-isotropic pressure coupling was applied, allowing independent scaling of the membrane plane and the perpendicular dimension. During the simulation, PBC box fluctuations exceeded the default strain limit. To allow for large volume changes, the maximum strain was increased from 0.1 to 0.35, enabling up to 35% box variation. This combined thermostat–barostat scheme ensures stable integration and proper sampling of the NPT ensemble under semi-isotropic pressure control, making it well-suited for simulations of membrane-containing systems. A 12 ns recommended trajectory output was selected for the production run. Two replicas of a total of 28 µs was collected for cholesterol binding analysis. Protein and cholesterol visualization and snapshot generation were performed using VMD 1.9.3 and UCSF ChimeraX [45, 46].

## 4.9 Data Availability

The dataset used in this study can be downloaded from t https://github.com/LynaLuo-Lab/CholBindNet.

## 4.10 Code Availability

Source code is available at https://github.com/LynaLuo-Lab/CholBindNet, which includes a docker for experimental reproducibility as well as https://github.com/LynaLuo-Lab/cholbindnet.github.io for a localhost website to test any .pdb file.

## Acknowledgements

Anton 3 computer time was provided by the Pittsburgh Supercomputing Center (PSC) through Grant 1R24GM154042 from the National Institutes of Health (NIH). The Anton 3 machine at PSC is made available by D. E. Shaw Research. This work was supported by NIH grants GM130834 (Y.L.L.), and the Calpoly Pomona research fund (S.C.K).

## 4.12 Author Contributions

A.H conducted model development and interpretation, A.B. conducted docking and MD simulations, I.R. carried out data curation and model comparison with existing methods. J.E.L helped A.H with final model codes for reproducibility. S.C.K and Y.L.L designed and supervised the project. All authors prepared the manuscript.

