## Supplementary Information for "CholBindNet: Interpretable Neural Networks for Cholesterol Binding Site Prediction"

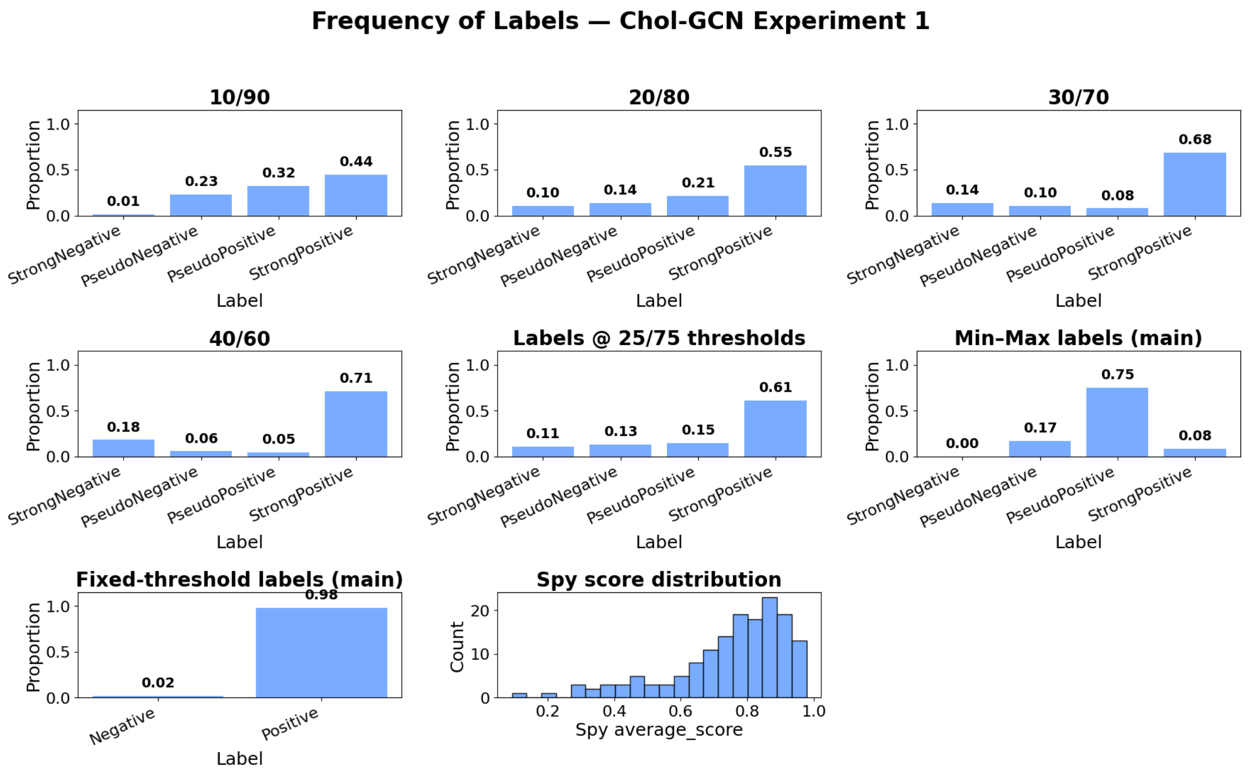


Figure S1: **Chol-GCN different threshold comparison.** Chol-GCN labeling of the test positive dataset under multiple thresholding strategies derived from spy probability distributions. The top and middle panels show percentile-based labeling schemes (10/90, 20/80, 30/70, 40/60, and the selected 25/75 thresholds), reporting the resulting proportions of samples assigned to negative, pseudo-negative, pseudo-positive, and positive classes. The min–max labeling scheme uses the minimum, mean, and maximum predicted probabilities as thresholds to define the four labeling classes. The fixed-threshold method assigns binary labels using a probability cutoff of 0.5, independent of the spy distribution. The bottom-right panel shows the distribution of spy average scores that motivates both the percentile-based and min–max thresholding strategies.

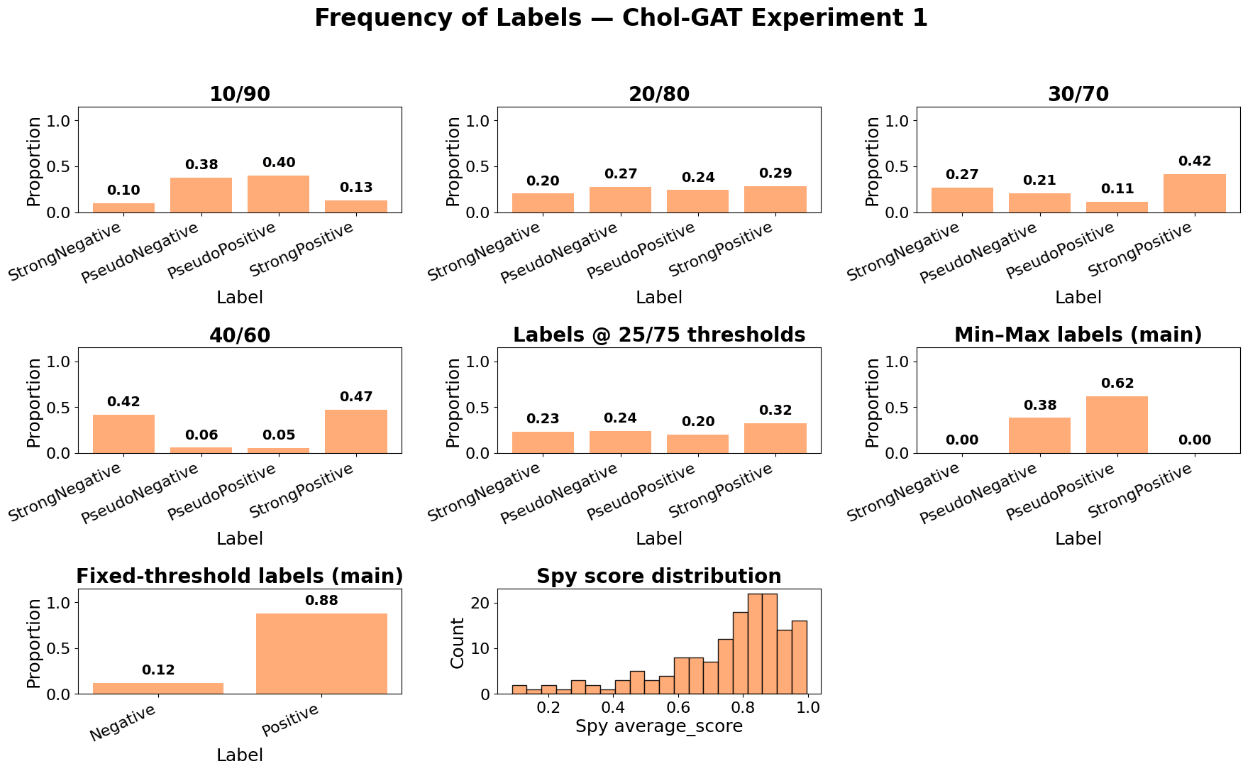


Figure S2: **Chol-GAT different threshold comparison.** Chol-GAT labeling of the test positive dataset under multiple thresholding strategies derived from spy probability distributions. The top and middle panels show percentile-based labeling schemes (10/90, 20/80, 30/70, 40/60, and the selected 25/75 thresholds), reporting the resulting proportions of samples assigned to negative, pseudo-negative, pseudo-positive, and positive classes. The min–max labeling scheme uses the minimum, mean, and maximum predicted probabilities as thresholds to define the four labeling classes. The fixed-threshold method assigns binary labels using a probability cutoff of 0.5, independent of the spy distribution. The bottom-right panel shows the distribution of spy average scores that motivates both the percentile-based and min–max thresholding strategies.

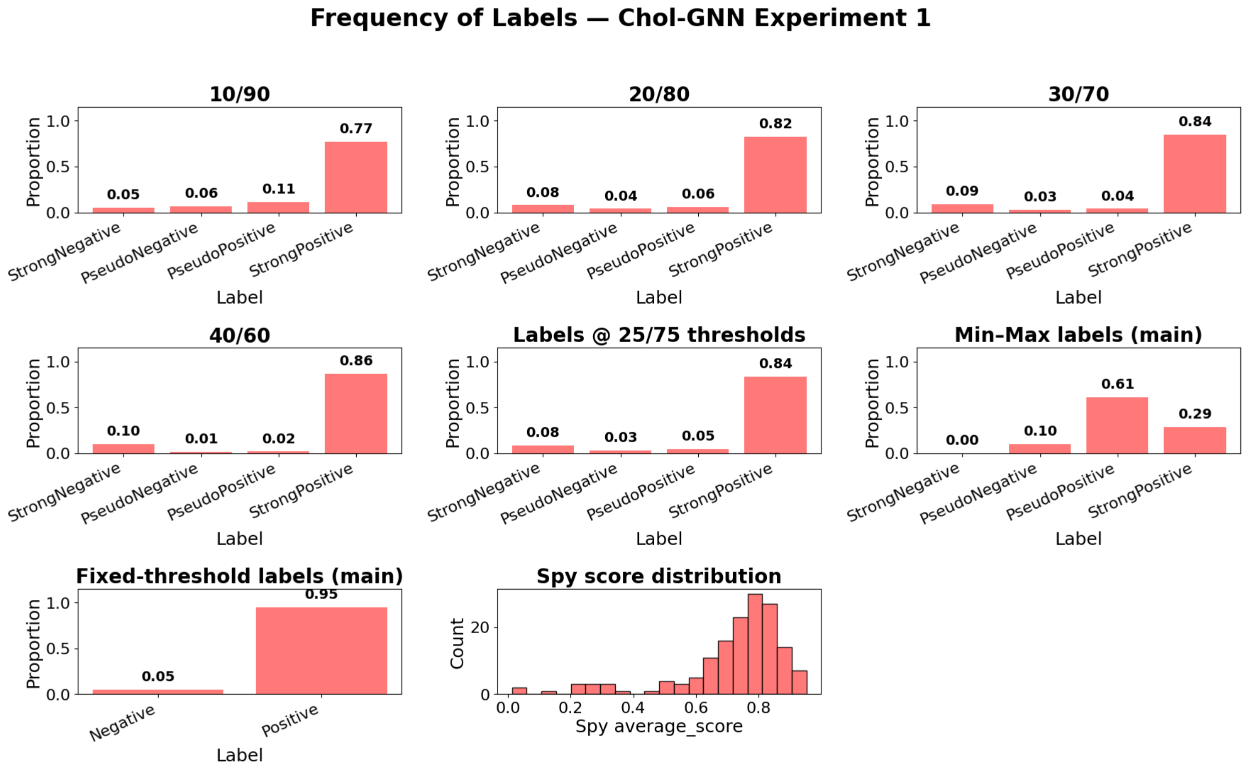


Figure S3: **Chol-GNN different threshold comparison.** Chol-GNN labeling of the test positive dataset under multiple thresholding strategies derived from spy probability distributions. The top and middle panels show percentile-based labeling schemes (10/90, 20/80, 30/70, 40/60, and the selected 25/75 thresholds), reporting the resulting proportions of samples assigned to negative, pseudo-negative, pseudo-positive, and positive classes. The min–max labeling scheme uses the minimum, mean, and maximum predicted probabilities as thresholds to define the four labeling classes. The fixed-threshold method assigns binary labels using a probability cutoff of 0.5, independent of the spy distribution. The bottom-right panel shows the distribution of spy average scores that motivates both the percentile-based and min–max thresholding strategies.

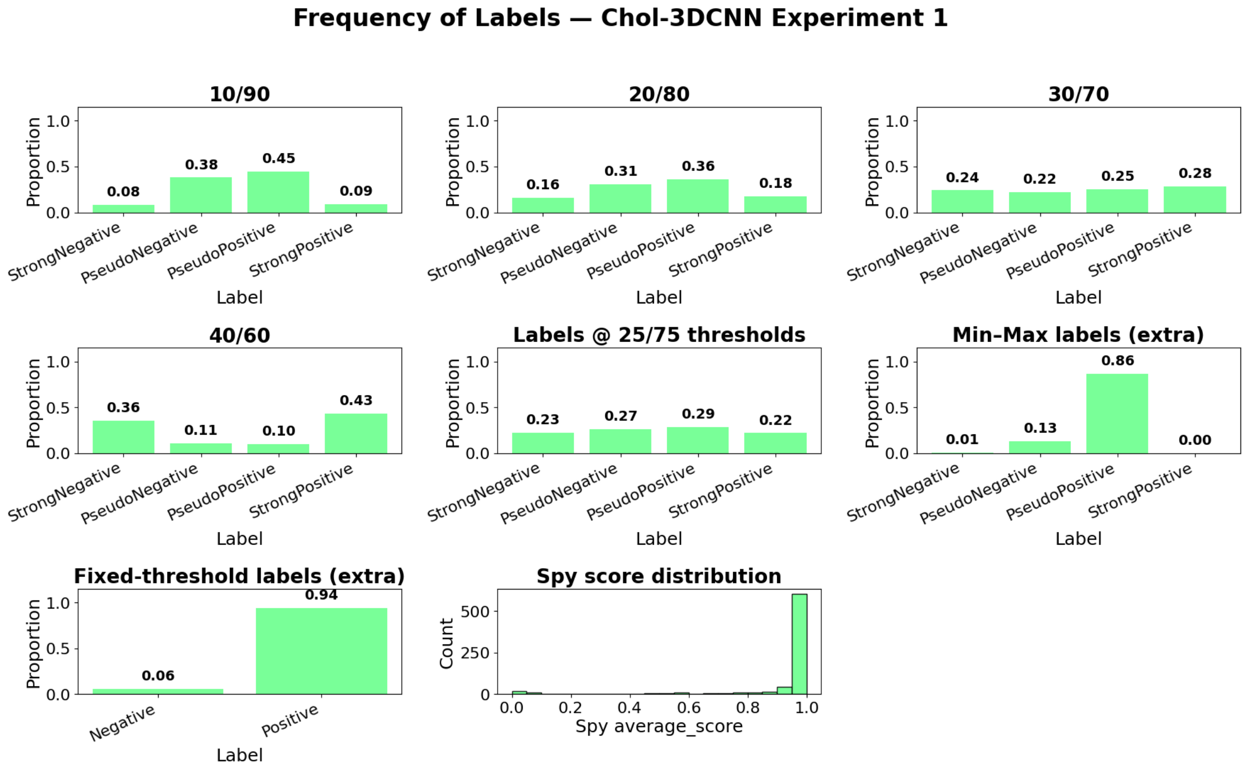


Figure S4: **Chol-3DCNN different threshold comparison.** Chol-3DCNN labeling of the test positive dataset under multiple thresholding strategies derived from spy probability distributions. The top and middle panels show percentile-based labeling schemes (10/90, 20/80, 30/70, 40/60, and the selected 25/75 thresholds), reporting the resulting proportions of samples assigned to negative, pseudo-negative, pseudo-positive, and positive classes. The min–max labeling scheme uses the minimum, mean, and maximum predicted probabilities as thresholds to define the four labeling classes. The fixed-threshold method assigns binary labels using a probability cutoff of 0.5, independent of the spy distribution. The bottom-right panel shows the distribution of spy average scores that motivates both the percentile-based and min–max thresholding strategies.

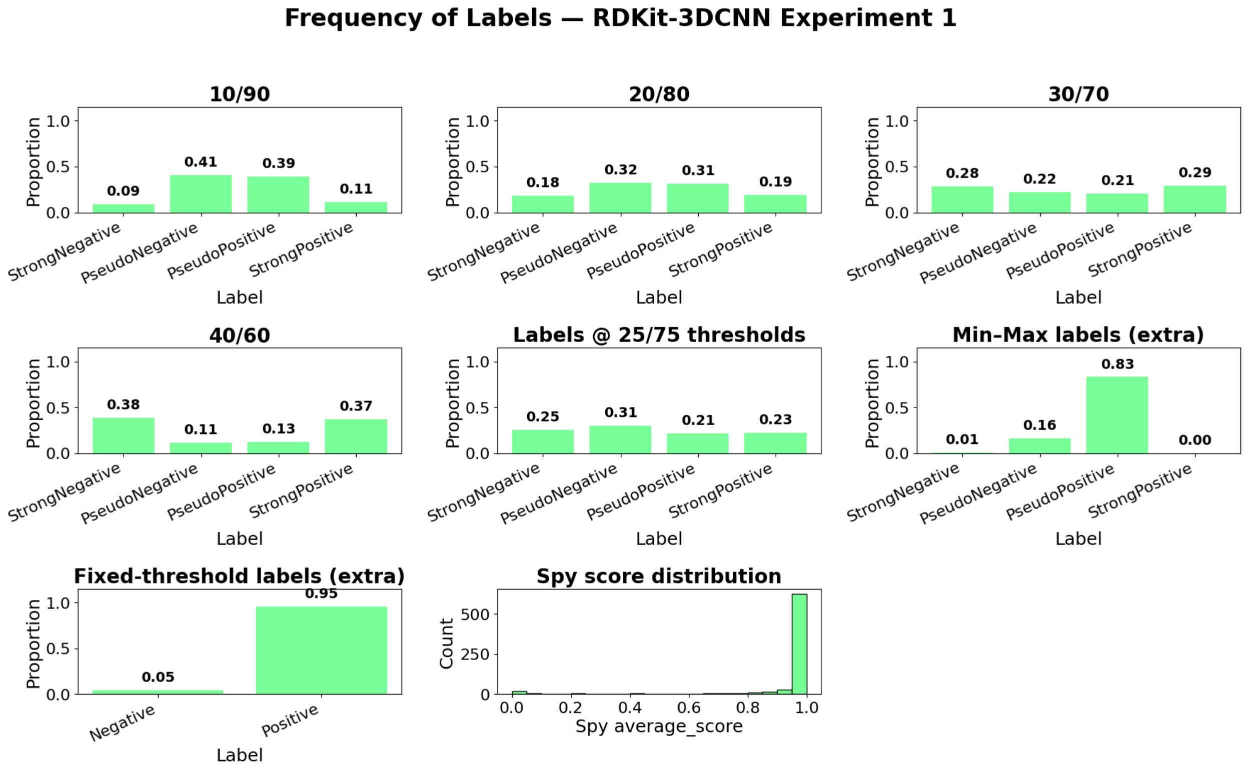


Figure S5: **RDKit-3DCNN different threshold comparison.** RDKit-3DCNN labeling of the test positive dataset under multiple thresholding strategies derived from spy probability distributions. The top and middle panels show percentile-based labeling schemes (10/90, 20/80, 30/70, 40/60, and the selected 25/75 thresholds), reporting the resulting proportions of samples assigned to negative, pseudo-negative, pseudo-positive, and positive classes. The min–max labeling scheme uses the minimum, mean, and maximum predicted probabilities as thresholds to define the four labeling classes. The fixed-threshold method assigns binary labels using a probability cutoff of 0.5, independent of the spy distribution. The bottom-right panel shows the distribution of spy average scores that motivates both the percentile-based and min–max thresholding strategies.

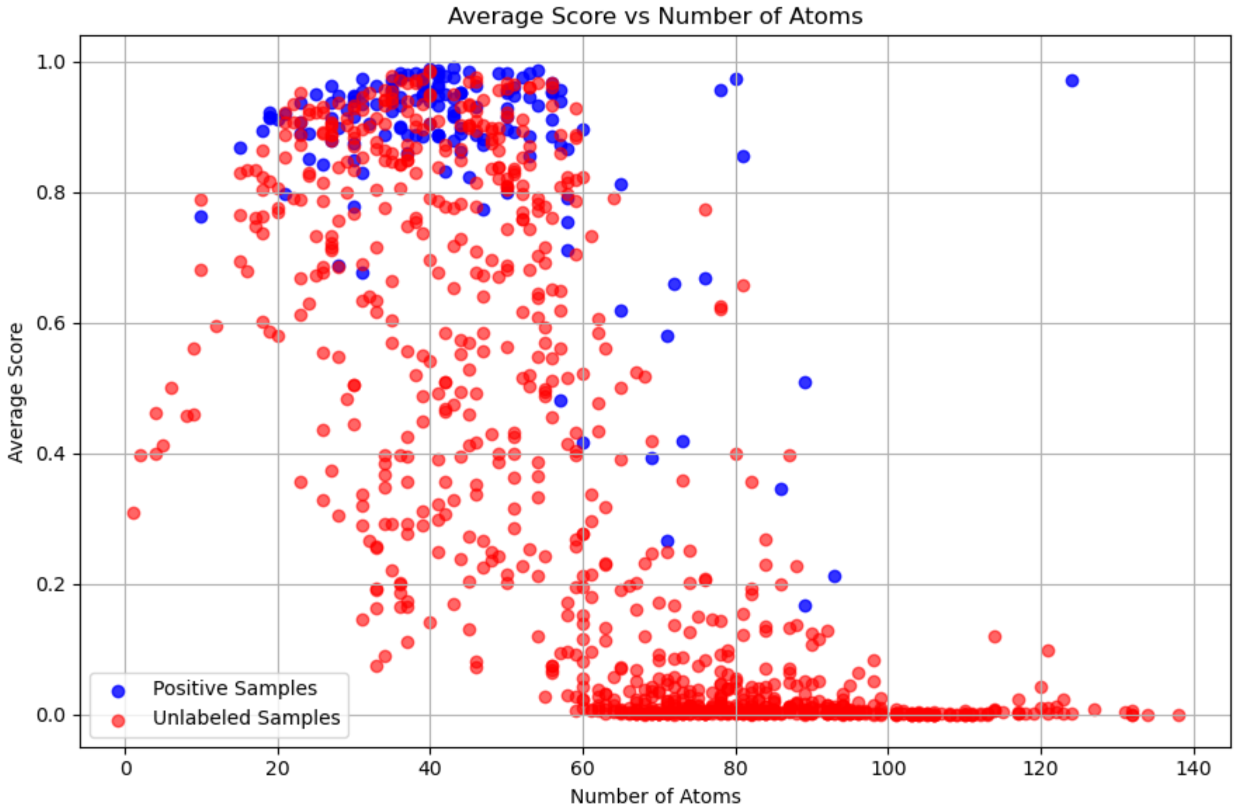


Figure S6: **Model probability score vs number of atoms in binding pocket.** Chol-GNN model predicted probability scores are plotted against the number of atoms in each binding pocket. Positive samples are shown in blue, while unlabeled samples are shown in red.

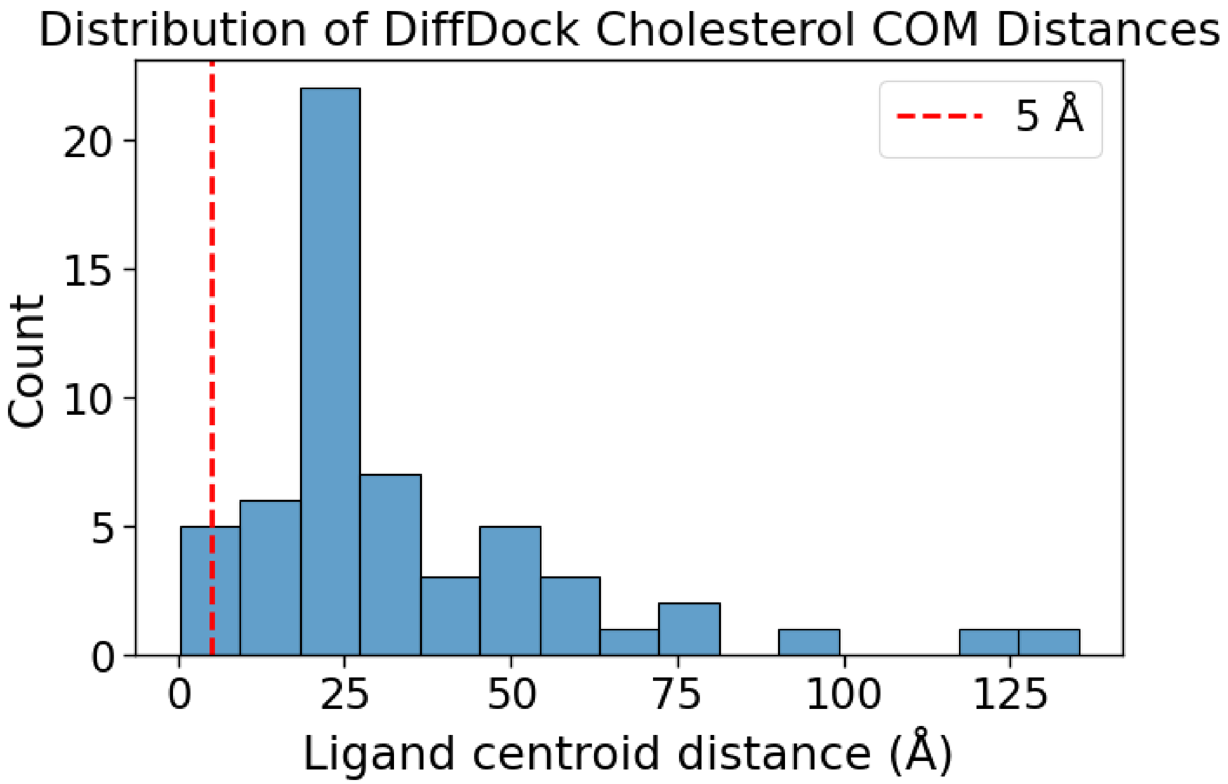


Figure S7: **The distribution of center-of-mass (COM) distances between the Diff-Dock predicted cholesterol and their corresponding native ligand positions.** This graph represents all 57 of the external validation dataset. The majority of predictions are found beyond 5 Å with only a small number below this threshold.

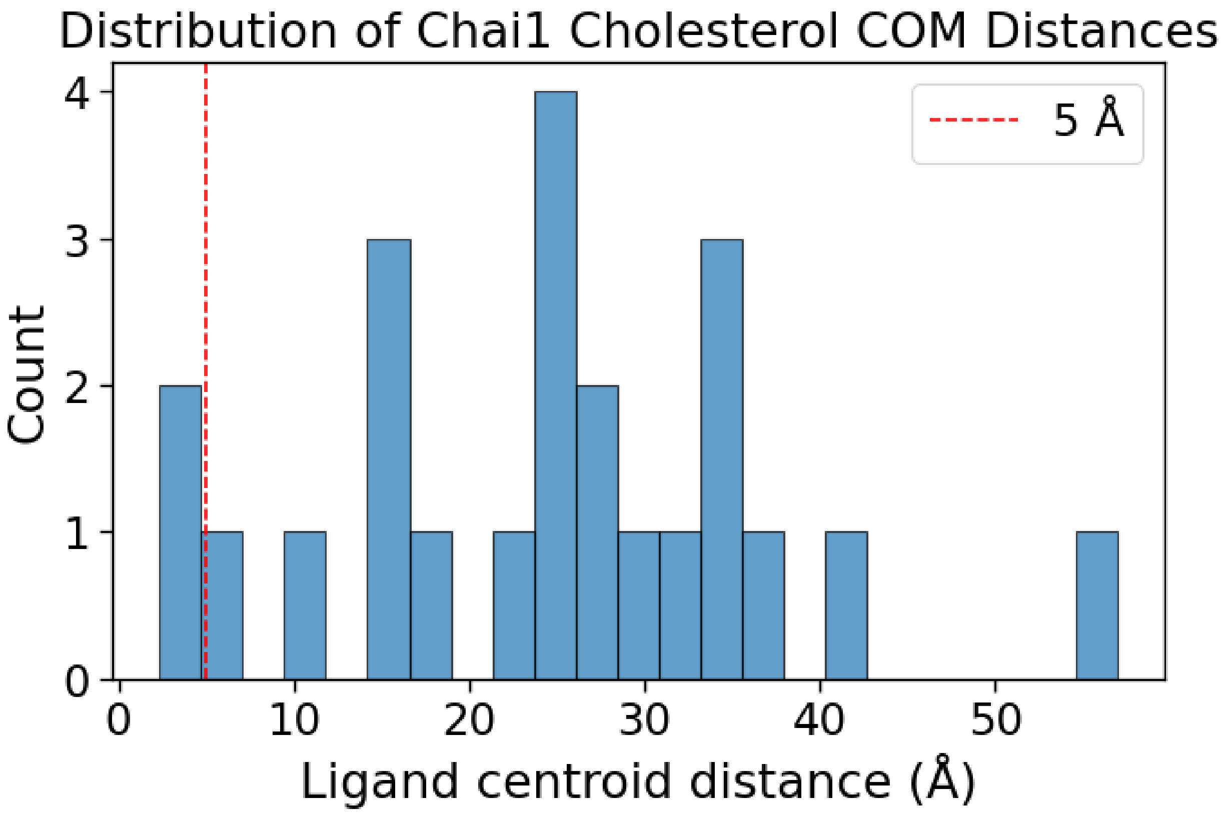


Figure S8: **The distribution of center-of-mass (COM) distances between the Chai-1 predicted cholesterol and their corresponding native ligand positions.** This graph represents 23 of the subset structures of the external validation dataset. Distances span a broad range, with the majority of predictions lying well beyond 5 Å . Only a small number of cases fall near or below this threshold, while most predictions show substantial displacement from the native binding site.

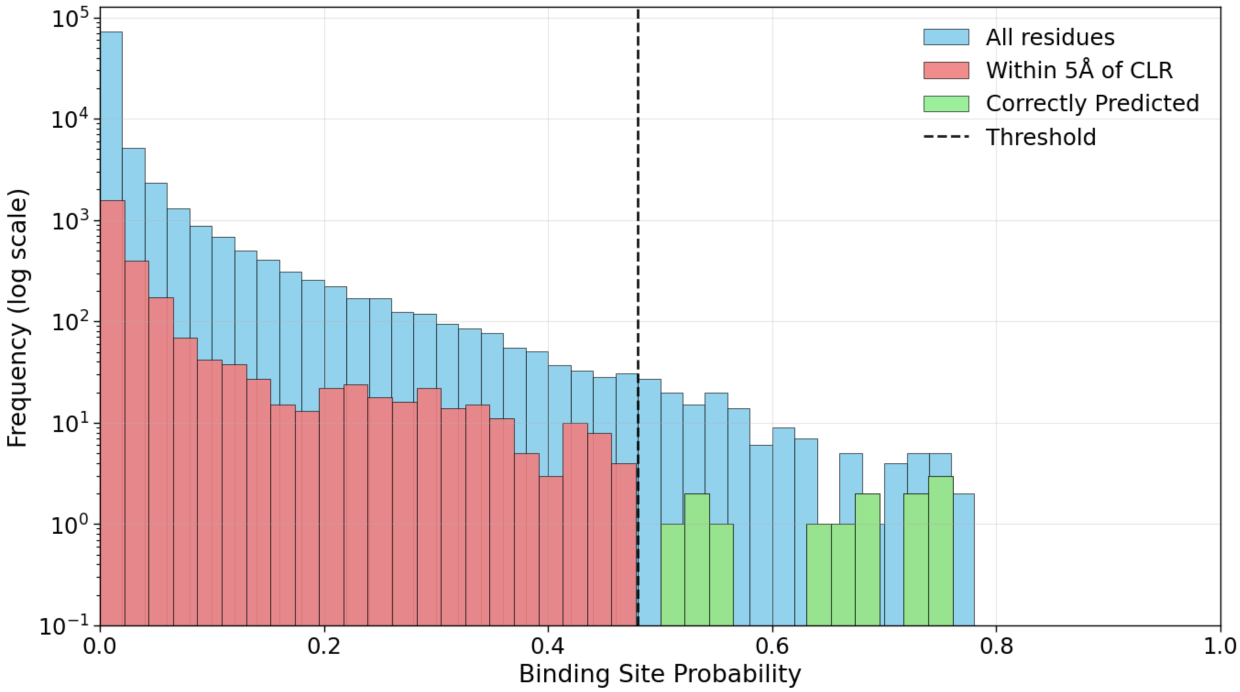


Figure S9: **LABind Predictions on CLR.** This image shows the distribution of binding site probability predictions generated by LABind. The blue bars represent all residues analyzed (a total of 85,678 residues), each assigned a binding site probability by the model. The red bars correspond to residues located within 5 Å of CLR in the experimental data. Using LABind's classification threshold of 0.48, the model identified only 13 residues, originating from four PDB structures, as high-probability binding sites.

Table S1: Encoded Atom Features 3DCNN Using RDKit

| **Columns** | **Feature Description** |
|---|---|
| 1–5 | Atom type (C, N, O, S, Other) |
| 6–8 | Hybridization type (SP, SP2, SP3) |
| 9 | Number of heavy atoms |
| 10 | Number of heteroatoms |
| 11 | Aromaticity indicator |
| 12–13 | Charge presence and charge sign (positive/negative) |
| 14 | Ring membership indicator |
| 15 | Acidic residues (ASP, GLU) |
| 16 | Basic residues (LYS, ARG) |
| 17 | Histidine (HIS) |
| 18 | Cysteine (CYS) |
| 19 | Polar uncharged residues (ASN, GLN, SER, THR) |
| 20 | Glycine (GLY) |
| 21 | Proline (PRO) |
| 22 | Aromatic residues (PHE, TYR, TRP) |
| 23 | Hydrophobic residues (ALA, ILE, LEU, MET, VAL) |
